# Neurons in auditory cortex integrate information within constrained temporal windows that are invariant to the stimulus context and information rate

**DOI:** 10.1101/2025.02.14.637944

**Authors:** Magdalena Sabat, Hortense Gouyette, Quentin Gaucher, Mateo López Espejo, Stephen V. David, Sam Norman-Haignere, Yves Boubenec

**Affiliations:** Laboratoire de Neurosciences Cognitives et Computationnelles, Département d’études cognitives, INSERM, Ecole Normale Supérieure, PSL University, Paris, France; Laboratoire des systèmes perceptifs, Département d’études cognitives, École normale supérieure, PSL University, Paris, France; Sorbonne Université, CNRS, Inserm, Institut de la Vision, F-75012 Paris, France; Oregon Hearing Research Center, Department of Otolaryngology-Head & Neck Surgery, Oregon Health & Science University. Portland, USA; Department of Biostatistics & Computational Biology, Department of Neuroscience, University of Rochester Medical Center, Rochester, USA; Department of Brain & Cognitive Sciences, Department of Biomedical Engineering, University of Rochester, Rochester, USA

**Author notes:** these authors equally contributed to the work (Sam Norman-Haignere, Yves Boubenec).

## Abstract

Much remains unknown about the computations that allow animals to flexibly integrate across multiple timescales in natural sounds. One key question is whether multiscale integration is accomplished by diverse populations of neurons, each of which integrates information within a constrained temporal window, or whether individual units effectively integrate across many different temporal scales depending on the information rate. Here, we show that responses from neurons throughout the ferret auditory cortex are nearly completely unaffected by sounds falling beyond a time-limited “integration window”. This window varies substantially across cells within the auditory cortex (∼15 to ∼150 ms), increasing substantially from primary to non-primary auditory cortex across all cortical layers, but is unaffected by the information rate of sound. These results indicate that multiscale computation is predominantly accomplished by diverse and hierarchically organized neural populations, each of which integrates information within a highly constrained temporal window.

## Introduction

Temporal integration is central to nearly all aspects of auditory perception. A key challenge is that information in natural sounds is distributed across a vast range of timescales. Speech and animal vocalizations, for example, are characterized by time-varying changes in pitch and spectrotemporal structure that unfold over tens of milliseconds, which are then assembled into longer sequences across hundreds of milliseconds or seconds^1^. Importantly, the rate at which this information varies is itself highly dynamic^2^. The rate of spectrotemporal changes, for example, is constantly varying, even within a vocalization, and the duration and density of vocalizations that compose a sequence are also highly variable. Thus, to derive information from complex sounds such as speech and animal vocalizations, the auditory system must be able to flexibly integrate across multiple timescales.

One hypothesis is that neurons in the auditory cortex can flexibly vary the time window over which they integrate information, depending on the rate at which information is varying. This hypothesis is motivated, in part, by prior work showing neurons in the auditory cortex perform flexible, context-dependent computations^3–5^. Artificial neural networks increasingly rely on flexible and nonlinear computational units whose temporal integration window can dynamically vary^6,7^, and such systems have shown promise as models of neural computation in the brain^8–13^. A second hypothesis is that individual neurons integrate across largely fixed timescales, but instantiate a diverse array of these integration scales across cells so as to support flexible multiscale integration across the neural population^14^. This hypothesis is motivated in part by prior work suggesting that cortical neurons show a diverse range of temporal scales even within a single cortical region ^15–18^.

Distinguishing between these hypotheses has been challenging, in part due to the difficulty of measuring integration windows from highly nonlinear systems such as the brain. Integration windows are often defined as the time window analyzed by a sensory system, and thus the window within which stimuli can alter the neural response^19–22^. Although this definition is simple and general, there have been no simple and general methods for estimating integration windows. Neurons in the auditory system are often modeled using spectrotemporal receptive fields (STRFs), which assume a fixed, linear mapping between a spectrogram and the neural response. STRFs, however, cannot model dynamic nonlinear processing, which is prominent in the cortex^23–26^, and likely plays an important role in flexible, multiscale integration^6,10,27–29^.

In this study, we leveraged a recently developed method for measuring integration windows from nonlinear (or linear) systems (the temporal context invariance or TCI method) (**Figure 1A**). The method directly tests the definition of an integration window by determining whether there is a stimulus duration for which the response is invariant to surrounding context stimuli. As a consequence, the method does not make any assumptions about the features that underlie the response (e.g., a spectrogram) or the relationship between those features and the neural response (e.g., linear or nonlinear). The TCI method was recently validated using artificial deep neural networks^7^ and macroelectrode recordings from the human auditory cortex^30^. Macroelectrodes, however, average activity across thousands of neurons within a local area^31–34^, which makes it impossible to determine if multiscale integration is accomplished by diverse populations of neurons integrating across largely fixed timescales or flexible integration within individual neurons.

**Figure 1.**
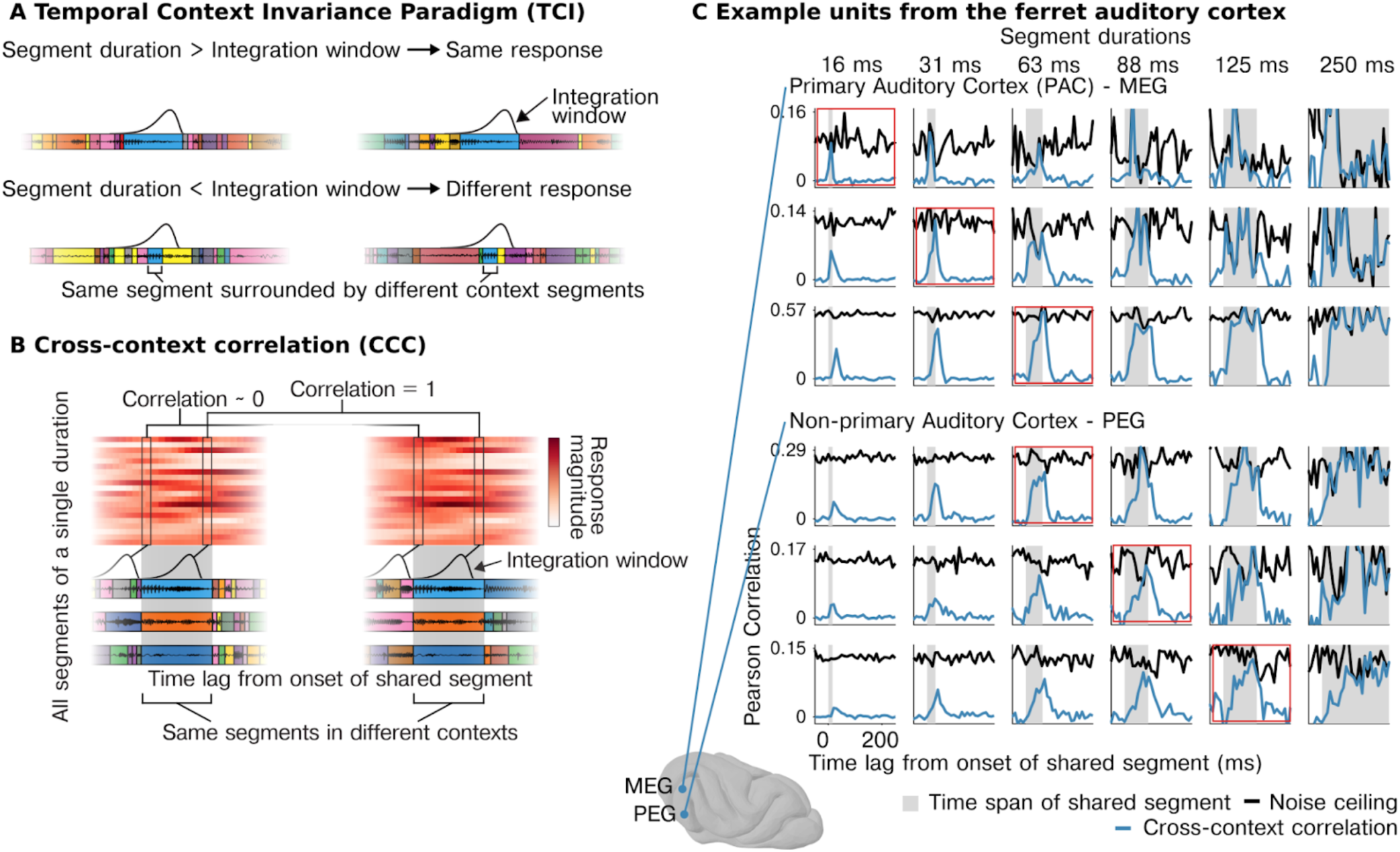
Neurons in auditory cortex show context-invariant responses. A, Segments of natural sounds were presented in different orders, leading to the same segment (blue) being surrounded by different “context segments”. **Top row**, If the neural response has an integration window that is less than the segment duration, there will be a moment when the window is fully contained within the shared segment, yielding the same response across both contexts. Waveforms for different segments are schematically indicated by colored boxes, and the integration window is plotted at the moment of maximum overlap between the two contexts. **Bottom row**, If the shared segments are shorter than the integration window, then the surrounding context segments can alter the response. **B**, Schematic of the “cross-context correlation” (CCC) analysis used to estimate context invariance. Response timecourses surrounding each segment are organized as a segment-by-time matrix (aligned to segment onset), separately for each context. We then correlate responses across segments between the two contexts for a given lag, which is equivalent to correlating corresponding columns (connected boxes) across matrices from different contexts (see text for details). A noise ceiling for the CCC is computed by performing the same analysis when the context is identical (using repeated presentations of each sequence). **C**, Example of the CCC (blue) and noise ceiling (black) from example multi-units in primary (MEG - *middle ectosylvian gyrus*, top) and non-primary (PEG - *posterior ectosylvian gyrus*, bottom) ferret auditory cortex. For all units, there is a lag and segment duration for which the CCC equals the noise ceiling, indicating a context-invariant response. The segment duration needed to achieve a context-invariant response varies substantially across units.

In this study, we applied the TCI method to microelectrode recordings from the ferret auditory cortex using ferret vocalizations, speech, and music across three experiments. We first tested whether cells in the auditory cortex showed evidence of a well-defined integration window, beyond which stimuli have little effect on the response, and if so, how this window varies across the neural population. We then examined whether integration windows reflect absolute time or vary with the information rate of sound by systematically manipulating the information rate using stretched and compressed sounds.

## Results

### Neurons in the auditory cortex integrate information within constrained temporal windows

We characterized the integration window of neurons in the auditory cortex of awake ferrets using the TCI method. We define the integration window as the time window within which stimuli can alter a neural response and beyond which stimuli have no effect. For example, an integration window of 100 ms would imply that the response to sound segments of 100 ms or longer will be unaffected by surrounding context segments. We note that while “context” has many meanings^35^, we operationally define context in our method as the surrounding segments.

In our method, we presented sequences of sound segments of varying durations in two different orders, such that each segment was surrounded by distinct “context” segments (**Figure 1A**). If the integration window is less than the segment duration, then there will be a moment when the integration window is fully contained within the segment, and thus produces the same response across the two different contexts (**Figure 1A**, top panel). In contrast, if the segment duration is shorter than the integration window, then the surrounding segments can alter the response (**Figure 1A**, bottom panel). Thus, if a neuron has a well-defined integration window, then the response to segments that are equal to or longer than the window will be context invariant.

We measured context invariance using the “cross-context correlation” (CCC) analysis. The analysis can be applied to any neural response timecourse. Here, the timecourse corresponds to the evoked spike rate to the segment sequences from our experiment. The first step of our analysis was to reorganize these timecourses as a matrix (**Figure 1B, Supplementary** Figure 1). Specifically, we extracted the response timecourses surrounding all of the segments of a given duration (e.g, 100 ms). We then compiled these timecourses as a segment-by-time matrix, aligned to segment onset. We compute a separate matrix for each of the two contexts using responses to the two different segment orderings (**Figure 1B, Supplementary** Figure 1A). Corresponding rows of these two matrices contain response timecourses to corresponding segments. These two matrices thus contain responses to many different “shared segments” surrounded by different context segments.

The cross-context correlation is then computed by correlating corresponding columns of these two matrices, where columns reflect a different time lag relative to the shared segment onset. At the start of the shared segment, the cross-context correlation should be approximately zero because the integration window will overlap the preceding segments, which are random between the contexts. As time progresses, the window will begin to overlap the shared segment, and the correlation will increase. If the window is less than the segment duration, there will be a time lag when the window is fully contained within the shared segment. At this time lag, the response will be the same between the two contexts, yielding a correlation of 1, modulo noise.

To correct for noise, we compute a noise ceiling by performing the same analysis but comparing responses to segments surrounded by the same context, using two repetitions of the same segment sequence (**Supplementary** Figure 1B). The noise ceiling thus provides an estimate of the maximum correlation possible for the CCC. In general, the reliability of the neural response will vary across units and across time/stimuli, since the response strength will vary across stimuli, which in turn affects the reliability. We therefore compute the noise ceiling separately for every unit, segment duration, and time lag, thus accounting for variation in response reliability across neurons and across stimuli. If neurons in the auditory cortex have a time-limited integration window, then there will be a segment duration and time lag for which the cross-context correlation reaches the noise ceiling.

We plot the CCC and noise ceiling for multi-units from different regions of the ferret auditory cortex (**Figure 1C**; see **Supplementary** Figure 2 for additional units). Strikingly, we found that for virtually all cells with a reliable response to sound, there was a segment duration and lag for which the CCC reached the noise ceiling, indicating a nearly fully context-invariant response. Once the CCC reached the noise ceiling, it tightly tracked the noise ceiling until segment offset, consistent with a temporally compact window composed of a single mode (i.e., not multimodal). These results indicate that nearly all sound-responsive cells in the auditory cortex show a temporally compact integration window beyond which stimuli have little effect on the neural response. Notably, the segment duration and lag needed to achieve a context-invariant response differed substantially across units and cortical fields (**Figure 1C**). For some units, segment durations as short as 15 ms were sufficient to yield a context-invariant response, while other units required longer segment durations of 32, 64, or 128 ms, particularly in non-primary regions of the auditory cortex (PEG).

We can also observe that for longer segment durations, the CCC tends to show a higher peak and longer temporal extent. The peak of the CCC reflects the extent of context invariance at the moment when the integration window maximally overlaps the shared segment. If the segment duration is longer, then the window will overlap the shared segment more and the context less. As a consequence, the peak will tend to be higher for longer segments. The temporal extent of the CCC reflects the time period during which the integration window is contained within the shared segment. If the segment duration is longer, the window will spend more time within the shared segment, extending the duration.

### Neural integration windows vary widely across the neural population, increasing in non-primary regions across all cortical layers

To quantify these effects, we used a computational model to estimate the integration window of each identified unit (345 units with reliable response to sounds across 3 animals, test-retest correlation r> 0.1; p<10^-5^ via permutation). The model was developed and has been validated in prior work^30^. The model parameterizes the integration window using a Gamma distribution, which is commonly used to model temporal windows^36,37^. The window is defined by three parameters: (1) the window width determines the temporal extent of the window and is defined as the smallest interval containing 75% of the mass (2) the window center determines the relative delay between the stimulus and the integration window and is defined as the window’s median (3) the window shape can vary from more exponential to more Gaussian. We varied these three parameters and found the window that best predicted the cross-context correlation. The prediction is computed by measuring the relative overlap of the window with the shared central segments compared with the surrounding segments. The corresponding equations are given in the Methods and have been derived and tested in prior work^30^. In practice, the window shape does not substantially impact the results (the predictions for different shapes do not vary substantially) and thus is not a focus of our analysis.

Figures 1-5 plot the estimated width of the model window (the smallest interval that contained 75% of the mass). The model also computes an estimated integration “center” (median of the parametric window), which can be thought of as the overall delay between the stimulus and the integration window. The center of the window was highly correlated with the window’s width and close to the minimal possible center given the window width, suggesting that neurons integrate information about as quickly as possible given the time window being analyzed (**Supplementary** Figure 3).

This analysis revealed a wide range of integration windows across cells spanning approximately 15 to ∼150ms (Figure 2A-D**; Supplementary** Figure 4 shows analogous results for integration centers). These integration times showed clear anatomical organization (Figure 2A): cells that were anatomically closer showed more similar windows (Figure 2B), and cells further from primary auditory cortex (PAC) showed longer windows (Figure 2C) (F_1,343_ = 84.79, *p* < 0.001, ^β^_distance to PAC_ ^= 0.35^ octaves/mm, CI = [0.275, 0.424]; N=345), leading to a substantial and significant difference between primary (*Median*=31.3 ms, *SD*=14.7 ms) and nonprimary auditory cortex (*Median*=54 ms, *SD*=11.3 ms) (Figure 2D) (F_1,343_ = 109.06, *p* < 0.001, β_region_ = −0.799 octaves, CI = [−0.949, −0.648]; N=345).

**Figure 2.**
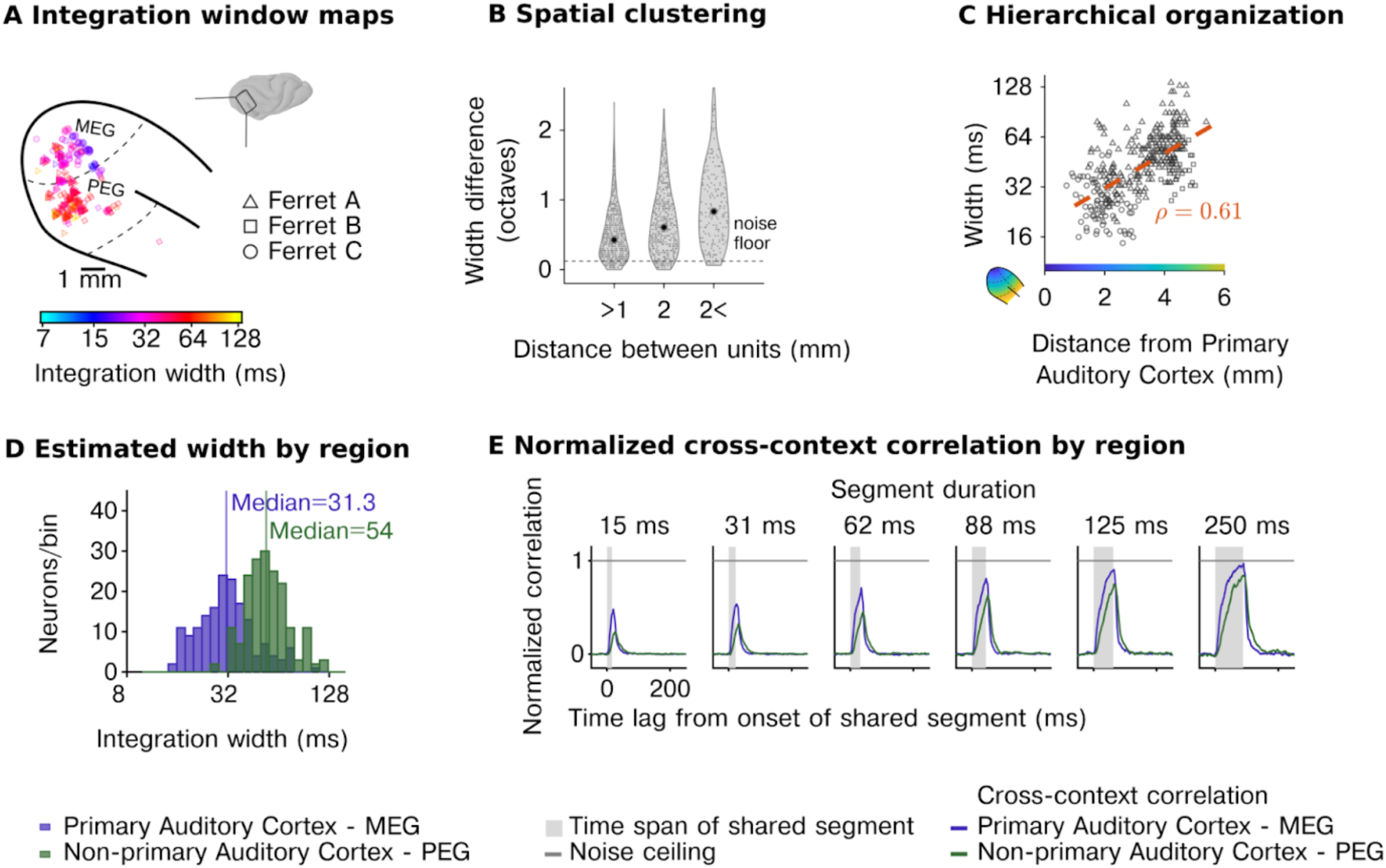
Diverse, hierarchically organized temporal integration windows across the neural population. **A**, Anatomical map of model-estimated integration windows in three animals (window width: smallest interval containing 75% of the window’s mass). **B**, The difference in integration windows between pairs of units as a function of their spatial distance, demonstrating that nearby units have more similar windows. **C**, Integration windows as a function of distance to the primary auditory cortex (see color map in inset). **D,** Histograms of integration windows for primary and non-primary auditory cortex showing substantial diversity across units and hierarchical organization. **E,** Median noise corrected cross-context correlation for all neurons in primary and non-primary auditory cortex. *Acronyms: MEG (middle ectosylvian gyrus) is a field of the primary auditory cortex. PEG (posterior ectosylvian gyrus) is a field of non-primary auditory cortex*.

To complement our model-based analyses, we also measured the average CCC in primary and non-primary auditory cortex and normalized this metric by the average noise ceiling, also computed separately for each region (Figure 2E). This analysis replicated our model-based results, with non-primary regions requiring substantially longer segment durations for the CCC to reach the noise ceiling. This analysis further demonstrated that even in non-primary regions, there is a time-limited window (∼150 ms), outside of which stimuli have very little effect on the neural response, suggesting that there is a compact integration window that strongly constrains the computations of virtually all neurons in the auditory cortex. We observed similar results for ferret vocalizations, speech, and music with substantially longer integration windows in non-primary regions for all three categories tested (**Supplementary** Figure 5).

Our first experiment shows that temporal integration is organized hierarchically across regions, but it did not enable us to investigate hierarchical organization between different cortical layers. To address this question, we repeated our experiments using laminar probes (231 cells with reliable response to sounds across 5 animals, test-retest correlation r> 0.1; *p*<10^-5^ via permutation) and grouped cells based on both their region and cortical layer (supragranular, granular, infragranular) using source density analysis (Figure 3A). We replicated our prior findings, showing a substantial increase in integration windows between primary and non-primary regions (F_1,21.47_ = 19.76, *p* < 0.001, β_region_ = −0.88 octaves, CI = [−1.270, −0.490]; N=231, Figure 3B). There was also a significant effect of layer (width: F_2,218.86_ = 6.87, *p* = 0.001) driven by shorter integration windows in the infragranular layers (β_infragranular_ = −0.486 octaves, CI = [−0.796, −0.175]). However, the magnitude of the effect was small both in absolute terms (ΔMedian_width_ = −4.73 ms) and compared with the effect of region (F_2,50.68_ = 30.88, *p <* 0.001). Consistent with these results, the average cross-context correlation was similar between layers but differed across regions, with non-primary areas requiring a longer segment duration and lags to reach the noise ceiling (Figure 3C). Thus, the dominant form of hierarchical anatomical organization appears to be region-based and not layer-driven.

**Figure 3.**
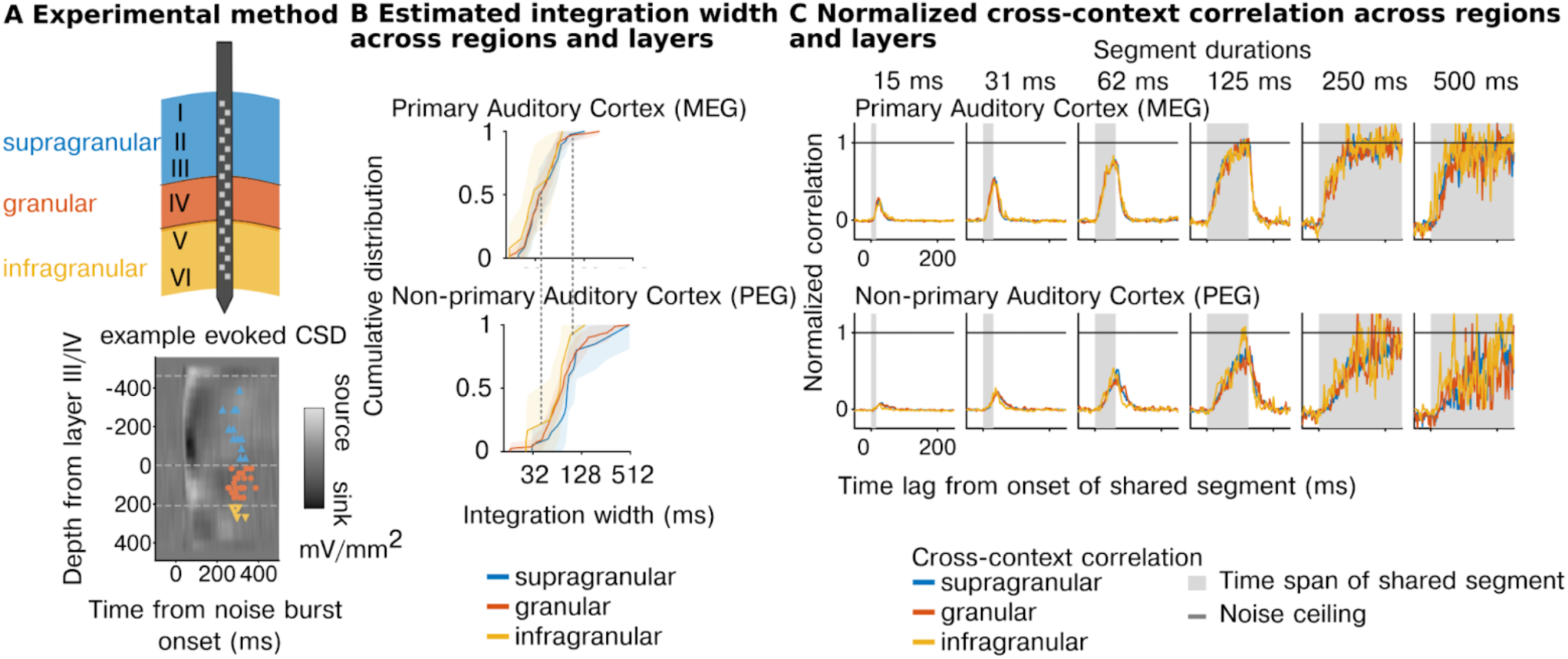
Integration windows show hierarchical organization across regions but not layers. A, *Top:* Schematic illustration of recordings from a laminar probe sampling supragranular (blue), granular (red), or infragranular (yellow). *Bottom:* Current Source Density (CSD) plot, evoked with a burst of noise for an example session. Individual markers correspond to recorded units and indicate approximate locations along the laminar probe. The color corresponds to the laminar classification of each unit. The colorbar reflects the spatial distribution of current flow in the extracellular space used to estimate the sources and sinks of change in measured potential along the laminar probe. Details of the laminar assignment are described in the method section. **B**, Cumulative distribution of estimated integration windows for primary (top) and non-primary (bottom) regions broken down by layer class (blue: supragranular, red: granular, yellow: infragranular). Dashed lines show corresponding points between plots from different regions so they can be visually compared. Integration windows are longer in non-primary regions but do not differ substantially between layers. **C,** Median noise corrected cross-context correlation separately for each layer (supragranular, granular, infragranular) and region (primary - MEG, nonprimary - PEG). *Acronyms: MEG (middle ectosylvian gyrus) is a field of the primary auditory cortex. PEG (posterior ectosylvian gyrus) is a field of non-primary auditory cortex.*

### Integration windows reflect absolute time and do not vary with the information rate of sound

Our analyses so far suggest that neurons in the auditory cortex integrate information within a highly constrained temporal window beyond which stimuli have little effect on the cortical response, and that this window varies substantially between neurons across the population. We next tested whether these integration windows reflect absolute time or instead vary with the information rate of the sound. To answer this question, we systematically altered the duration and rate at which sound information varies by stretching and compressing sounds by a factor of (preserving frequency/pitch) (Figure 4A). If the neural integration window varies with information rate (rate-yoked integration), the window should appear to scale with the magnitude of stretching and compression. In contrast, if the integration window reflects absolute time (time-yoked integration), the window will be unchanged by this manipulation. We tested both speech and ferret vocalization because they contain many different forms of time-varying structure that are ecologically relevant to the animal (particularly for ferret vocalizations). We focused our experiments on a non-primary region (PEG) because we expected rate-yoked integration, if present, to be strongest in higher-order regions ^16,38^.

**Figure 4.**
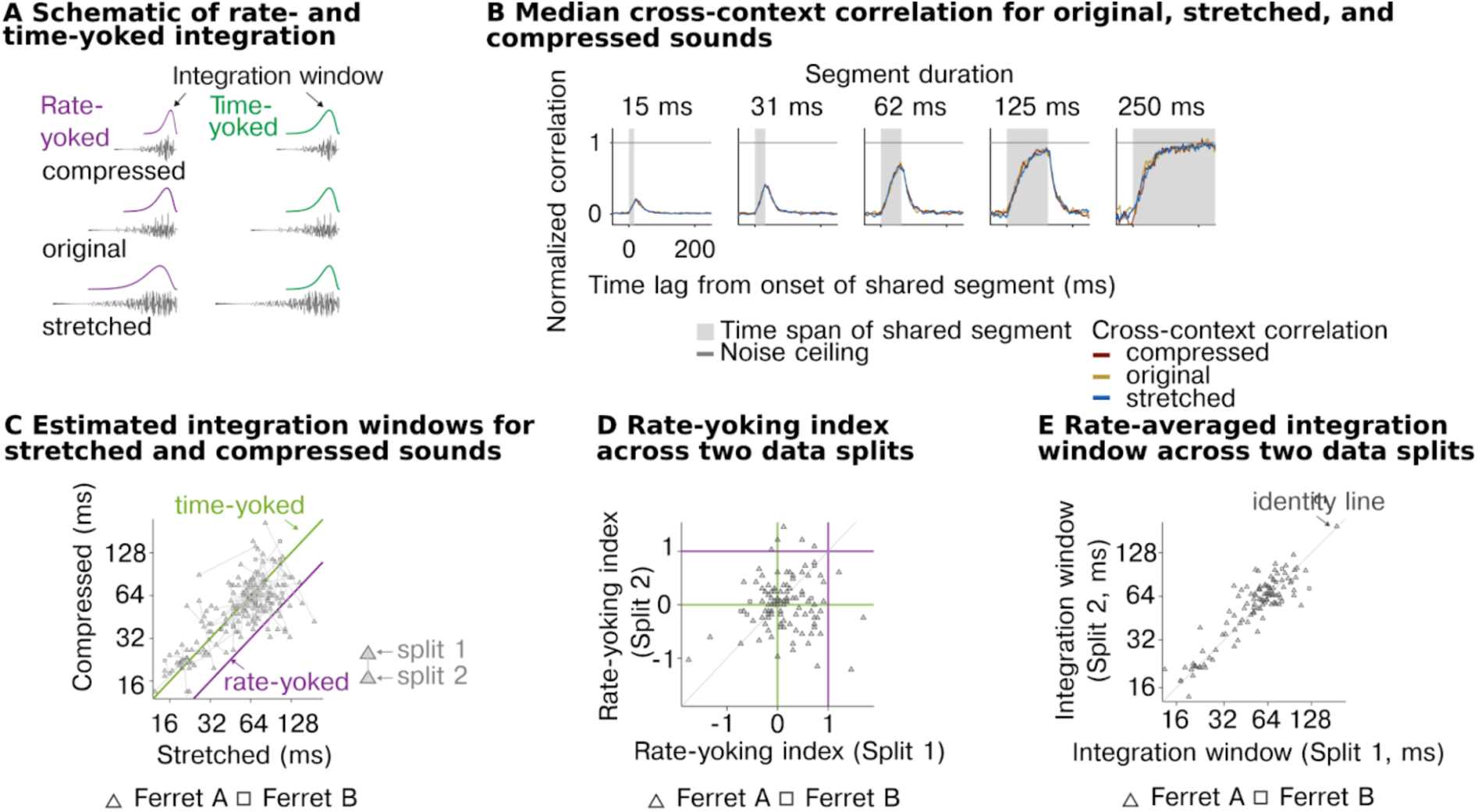
Temporal integration is invariant to information rate. **A**, Schematic illustration of the predictions for time-vs. rate-yoked integration. Stretching and compression will expand and shrink the integration window if it is yoked to the information rate of sound (purple, left) but not if it reflects absolute time (green, right). **B**, Median normalized CCC across all recorded units for stretched, original, and compressed stimuli. **C**, Integration windows for all units for compressed (x-axis) and stretched (y-axis) stimuli. Green and purple lines show the prediction from a time-yoked and rate-yoked response. **D,** Reliability of rate-yoking index across two independent data splits. **E,** Reliability of integration windows averaged across stimuli rates for comparison.

**Figure 5.**
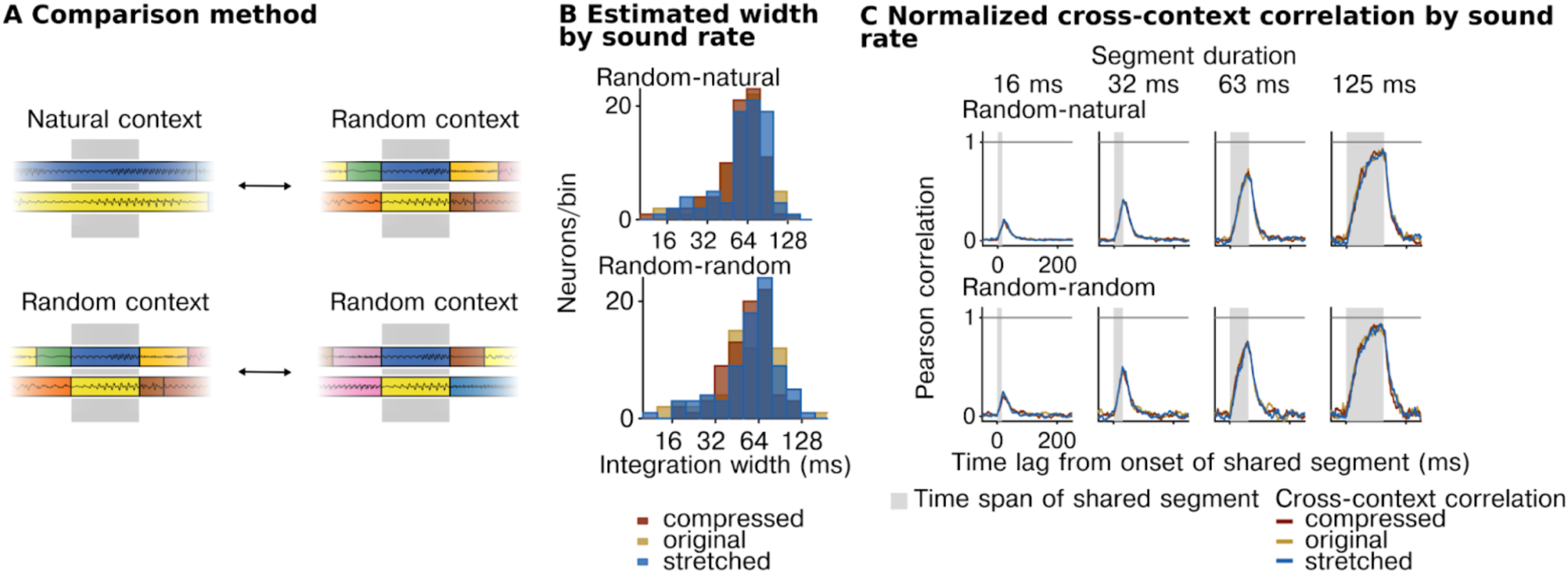
Integration windows estimated using natural context. **A**, We measured responses to segments that were subsets of longer segments and thus surrounded by their natural context (top, left) as well as segments surrounded by randomly selected segments. This figure plots results computed by comparing responses to natural and random contexts (top row of B&C), as well as results when comparing two different random contexts (bottom row of B&C). **B,** Histograms of estimated integration windows for different sound rates. **C,** Normalized cross-context correlation for different sound rates.

As a simple summary metric, we first measured the average cross-context correlation for all units from the non-primary auditory cortex for each of the different rates, pooling data from both sound categories (Figure 4B). We found that the average CCC was virtually identical across the three rates for all segment durations tested, suggesting integration windows reflect absolute time and do not vary with the information rate (see **Supplementary** Figure 6 for more example units). We next quantitatively compared the integration windows for different stimulus rates, separately estimated for each cell using our computational model (Figure 4C). If the integration windows are fully rate-yoked, integration windows for stretched sounds should be twice as long on average as the integration windows for compressed sounds, yielding an octave shift on a logarithmic scale (Figure 4C, purple line). If the integration windows are time-yoked, the windows should cluster around the line of unity (Figure 4C, green line). We found that the estimated windows clearly clustered around the line of unity, consistent with our previous analysis suggesting time-yoked integration dominates. We observed clear evidence of time-yoked integration for both sound categories tested (speech and ferret vocalizations) (**Supplementary** Figure 7).

There was substantial variation from cell to cell in the estimated integration window for stretched and compressed speech (Figure 4C). To test if this variation reflected genuine variation in rate yoking or just variation due to unreliable sources of noise, we measured the integration window for each cell and rate using two independent splits of data. We then computed a measure of rate yoking separately for each data split, and correlated this measure across units between the two splits (Figure 4D). Rate yoking was measured by subtracting the integration window for stretched and compressed speech on a logarithmic scale and dividing by the 2-octave difference in rates. The metric thus provides a simple, graded measure of the extent of time-yoked (0) vs. rate-yoked integration (1) (note that due to noise, the metric can exceed the 0 and 1 bounds). The effect of rate of the stimuli was marginally significant (width: F_2,664.58_ = 2.53, *p* = 0.08, β_stretched_ = 0.16 octaves, CI = [0.019, 0.308], β_compressed_ = 0.105 octaves, CI = [-0.039, 0.249]; N = 702 (observations for 117 neurons)), however, we found that the variation from cell to cell in the strength of rate yoking was not reliable across data splits (Spearman rank correlation of 0.09, p = 0.52 via bootstrapping), indicating that it mostly reflected unreliable source of noise in the data (Figure 4D). In contrast, the overall integration window, averaged across rates, was highly reliable (Spearman rank correlation of 0.72, p < 0.0001 via bootstrapping) (Figure 4E).

In our primary analysis, we pooled across two different types of context comparisons. The simplest comparison is the one illustrated in Figure 1A, where we compare responses to the same segment surrounded by randomly selected segments. Because they are randomly selected, these segments differ from those that would typically surround a segment in natural sounds. Thus, it is possible that the reason we observe time-yoked integration is that our analysis is not sensitive to natural contextual processing. To address this possibility, we therefore performed a second analysis to test whether similar results would hold when investigating a natural context. This analysis was enabled by the fact that shorter segments were created by subdividing longer segments. As a consequence, there were many cases in which a segment was surrounded by its natural context as a part of a longer segment. We could therefore measure and compare responses to two different types of context: (1) random context, where a segment is surrounded by unrelated segments, and (2) natural context, where the shared segment is a part of a longer one, preserving its original temporal surroundings (visual schematic Figure 5A). When computing the cross-context correlation, one context must be random so that the two contexts differ, but the other can be natural (Figure 5A, top row) or random (Figure 5A, bottom row). Critically, the only way to observe a context invariant response when comparing natural and random contexts is for the response to be invariant to the natural context. The natural-random comparisons provide a strong test of the extent of naturalistic contextual processing and thus arguably a stronger test of the rate yoking hypothesis.

In practice, we observed very similar results for both types of context comparisons (Figure 5 **B&C**). In particular, we found that the cross-context correlation was very similar for natural, stretched, and compressed speech, even when measured using natural context comparisons, indicating time-yoked integration. In addition, the fact that our results are similar for the two types of context comparisons indicates that all of our earlier results hold even using an analysis that is sensitive to naturalistic contextual effects.

## Discussion

Our study demonstrates that neurons in the auditory cortex integrate information across a time-limited window beyond which sounds have little influence on the cortical response. This window varies by approximately 3 octaves (∼15 to 150 ms) across cells in the auditory cortex and exhibits clear anatomical organization, with two to three times longer integration windows in non-primary regions and little difference within a cortical column. In contrast, these windows do not vary with the information rate and duration of sound structures (e.g., vocalizations), suggesting that they are a fundamental property of the auditory system and not a property of the sounds analyzed by the auditory system. These results indicate that multiscale computation is accomplished by diverse, hierarchically organized populations of neurons, each of which integrates information across a specific, time-limited, and context-invariant temporal window.

### Implications for models of multiscale computation

Understanding the computations that enable the auditory system to integrate across the multiscale temporal structure of natural sounds is an important goal of auditory neuroscience. Multiscale integration is computationally challenging in part because information in sound is organized across many different timescales, and these timescales are highly variable and context-dependent.

Our results show that virtually all neurons in the auditory cortex integrate information with a compact, temporal window beyond which stimuli have little effect on the neural response. This finding is highly nontrivial, since highly nonlinear, recurrent systems such as the brain are capable of integrating across potentially very long timescales^7,39^. Our results suggest that this window is temporally compact, since multi-peaked windows would produce oscillations in the cross-context correlation, which were not observed empirically. We also find that the center of the window is close to the minimum possible for a given integration width (**Supplementary** Figure 3). For a causal system whose responses depend on the past, the integration width, which reflects the total analysis time, places a constraint on how quickly information can be processed. If a neural system is processing information quickly, there will be no temporal gap between the stimulus and the window. As a consequence, the integration center will be close to the minimum possible center given the integration time, which we empirically find to be the case.

Our second key result is that integration windows show substantial heterogeneity across the neural population, with nearly 3 octaves of variation. We show that this variation is hierarchically organized across regions, but not across cortical layers. We note that increasing integration windows are not an inevitable consequence of hierarchical organization (as evidenced by the weak effect of layer). For example, hierarchical recurrent neural networks can show similar integration windows across their hierarchy^7^, and training RNNs on challenging auditory tasks does not necessarily induce increasing integration windows across model layers^40^. We note that while we observe a substantial increase in integration windows between regions, there is still substantial heterogeneity in integration windows within a region that exhibits anatomical clustering, suggesting that within a region, the auditory cortex performs multiscale temporal analysis using spatially organized neural populations.

Critically, we provide evidence that the temporal windows of neurons in the auditory cortex do not vary with the rate at which information varies in natural sounds. In particular, we find that temporal windows, on average, are completely unaffected by stretching and compression, even though stretching and compression change the rate at which virtually all acoustic structure varies, as well as the duration of all sound events (e.g., vocalizations). This result is also highly nontrivial. Nonlinear systems are highly capable of flexibly varying their integration window, and our TCI method is sensitive to these types of changes and can robustly detect rate-yoked integration when present^7^. Collectively, these results demonstrate that multiscale computation is performed by diverse populations of neurons, each of which integrates information within a highly constrained temporal window beyond which stimuli have very little effect on the neural response and which is invariant to the information. These results suggest that flexible temporal integration is accomplished by flexibility at the neural population level rather than flexibility within single neurons. Dean et al., (2008)^41^ found that neurons in the inferior colliculus adapt to changes in sound level statistics, and that the adaptation time did not vary with the temporal rate at which level changes occurred. Sound level, however, is only one feature of the sound, and adaptation to sound level is only one aspect of the neural response. Integration windows reflect the effects of all stimulus processing in natural sounds. Our findings thus suggest a much more general conclusion that the total amount of time being analyzed by the auditory cortex at any given moment does not vary with the rate at which information changes in natural sounds.

Previous research has observed that arousal and cognitive tasks can alter neural receptive fields (e.g., frequency tuning) even in the primary auditory cortex^16,42,43^, but has not investigated whether task demands alter integration windows. Our study shows that temporal integration is sensitive to the stimulus information rate, but it did not investigate the impact of attention or cognitive state. Future research could investigate whether cognitive variables such as attention or arousal modulate neural integration windows by using the TCI method to quantify integration windows for different tasks and state-dependent variables (e.g., pupil).

Our finding of hierarchical organization is consistent with a prior study that also observed hierarchical organization in the human auditory cortex^30^. This prior study used macroelectrodes, which pool activity from thousands of neurons and thus was not able to test whether individual neurons show a time-limited integration window, characterize the heterogeneity across the neural population, or examine layer-wise effects. Prior work has found that the functional organization of the human auditory cortex differs substantially from that of ferrets^44^, likely in part due to the unique importance of speech and music to human hearing, and thus a priori, it was entirely possible that we might have observed a very different organization in ferrets. Our findings, therefore, provide novel evidence that region-based hierarchical integration, in which longer timescale computations are performed using the output of shorter timescale computations, is a fundamental property of biological auditory systems that is not limited to the human auditory cortex.

The integration windows observed in this study are shorter than those observed in the human auditory cortex, which range from about 50 to 400 ms. This difference could reflect differences in the neural measures employed (e.g., macro-vs. micro-electrodes) or could reflect species differences. For example, prior work shows that neural populations with longer integration windows (< 200 ms) in the human auditory cortex show strong selectivity for speech and music^30^, which is absent from ferret auditory cortex^44^, suggesting that these longer timescales may have evolved or developed to process speech-or music-specific structure.

Prior studies have shown that the response latency and response duration increase across the cortical hierarchy in the ferret cortex^38,45^. These studies provide evidence for hierarchical temporal processing and thus are broadly consistent with our findings. Response delay and response duration, however, are distinct from the neural integration window. For example, two neurons with the same integration window can exhibit very different response delays. Similarly, a longer response duration does not necessarily imply a longer temporal integration window, as a system can retain activity or memory traces without integrating new information (e.g., Sussillo & Barak, 2013^46^). Nonetheless, these different measures of temporal processing may empirically be correlated. For example, we find that the center of the integration window is close to the minimum possible center, given the integration width, which suggests that the integration time will be correlated with the response delay. This observation, however, is an empirical result, not an inevitable consequence, of our measures, and suggests that information is integrated about as quickly as possible given the time being processed.

### Relationships to other types of neural timescales

One approach to characterizing the sensory responses is to attempt to learn an explicit functional mapping (or “encoding model”) between the stimulus and the neural response. Neural responses in the auditory system have been classically modeled using spectrotemporal receptive fields, which fit a linear mapping between a time-frequency spectrogram-like representation and the neural response^47^. STRFs, however, are unable to predict much of the neural response variance in the auditory cortex, particularly for natural sounds, due to prominent nonlinearities between the spectrogram and the neural response^48^ that are not captured by the model^6,49^. Some prior studies have fit more complex encoding models, such as deep neural networks^6,23,50^. Rahman et al., (2019)^50^, for example, included an exponential decay function as an ingredient of their deep learning model and showed that the time constant of this exponential varied substantially across neurons in the auditory cortex, varying from as little as 5ms up to 500 ms. A benefit of this approach is that it can be applied to completely natural stimuli without the need for reordered sound segments. A benefit of the TCI method is that it does not make any assumptions about the features that underlie the response (e.g., spectrogram) or how these features are mapped to the neural response (i.e., exponential decay function), and thus provides a more general measure of the integration time. While absolute integration times from the two approaches are not directly comparable, applying the TCI method to the predictions of models, such as those used by Rahman et al., would provide a useful means to test whether a model can replicate the neural integration time.

An alternative approach that does not require fitting a complete encoding model is to simply measure temporal characteristics of the neural response to a sound. For example, prior studies have measured the rate of temporal modulations in the neural response^51^, which has revealed that the upper frequency limit at which the neural response will “phase-lock” to a stimulus frequency slows from the periphery to the cortex. Phase locking, however, does not provide a measure of the neural integration window. For example, fast neural modulations could be produced by a neuron that integrates information across either a long or short window (e.g., integrating across many cycles of an oscillation or just a single cycle). Other studies have attempted to measure the time period in the neural response that contains information about a stimulus or stimulus property (sometimes referred to as the “encoding window”)^22^. While useful, this measure is highly stimulus-dependent and will vary with the duration and information rate of sounds. Other studies have measured the extent to which the neural response timecourse expands or contracts as the stimulus is stretched or compressed^52^. However, the neural response will tend to expand and contract with the stimulus, even if the integration window remains fixed, and this metric thus does not provide a good estimate of the integration window or whether this window varies with the information rate. By contrast, the TCI method can directly estimate the neural integration window, independent of many stimulus properties such as the information rate and duration of sounds, as we show here.

Recent research has characterized the “intrinsic timescales” of neural systems, often by measuring the autocorrelation of the neural response after removing stimulus-driven changes^53–57^. Intrinsic timescales are distinct from integration windows, since they do not directly specify anything about the relationship between the stimulus and the neural response. In contrast, integration windows are fundamentally defined with respect to the stimulus. Empirically, however, intrinsic timescales have been shown to increase across the cortical hierarchy, potentially analogous to the hierarchical organization that we observe here. Thus, one possibility is that there are shared underlying neurobiological mechanisms (e.g., changes in synaptic integration time constants) that couple intrinsic timescales and stimulus integration windows, and cause both to increase across the cortical hierarchy. Future research could explore this link by applying the methods used here to computational models of intrinsic timescales (i.e., Zeraati et al., (2022)^58^), which is feasible because our method is effective in nonlinear models.

### Methodological choices, limitations, and future directions

Many prior studies have demonstrated stimulus-driven effects that extend beyond the integration timescales observed in this study. For example, stimulus-specific adaptation effects have been observed over multi-second timescales^18,29,59,60^, and effects of learning and memory can be observed across minutes, hours, and days^16,61^. Thus, there are likely multiple timescales over which stimuli can influence the neural response, in part reflecting different neurobiological mechanisms (e.g., adaptation, learning, memory, etc.). Our results demonstrate that the cumulative effects of these longer timescale processes are small during the processing of natural sounds compared with the effects of stimulus variation within the window. Indeed, we find that for virtually every unit in the auditory cortex, there is a time period from approximately 15 to 150 milliseconds, where the cross-context correlation closely tracks the noise ceiling, indicating that surrounding stimuli have virtually no effect on the neural response. One caveat is that the noise ceiling measured by the TCI method will reflect all responses that are reliable across a long sequence of sounds. Thus, any neural responses that change between two presentations of the same sequence will be treated as noise. As a consequence, if there were learning and memory processes that took place across repetitions of an entire sequence^62^, our method would not be sensitive to them.

We used uniform stretching and compression (preserving pitch) because this alters the information rate and duration of virtually all sound properties by a known magnitude, which is useful because we do not know which features are important to the neural response. Recently, we showed that this approach is able to reveal a near complete transition from time-yoked to rate-yoked information across the layers of a highly nonlinear deep artificial neural network (DANN) trained on challenging speech recognition tasks, even though the models were only ever exposed to natural speech without any compression or stretching. This finding demonstrates that our method can cleanly detect rate-yoked integration, even from a highly complex nonlinear system, where the features and structures that underlie the system’s response are unknown. Nonetheless, a potential limitation of our approach is that uniform stretching and compression are not entirely natural, and it is possible that biological systems might show greater evidence of rate-yoked computation for more natural rate manipulations.

Our study focused on characterizing integration windows in the auditory cortex. We find that the auditory cortex integrates across a multiplicity of timescales spanning more than 3 octaves, but that these timescales are nonetheless relatively short, never exceeding approximately 150 ms. An important question for future research is how integration windows are structured beyond the auditory cortex in higher-order cognitive and motor regions, as well as in associative multisensory regions. Prior research suggests that neural timescales increase beyond the auditory cortex^55,56,63^, and it is thus possible that stimulus-driven integration windows show longer multisecond timescales in these regions. Another possibility is that integration windows become increasingly flexible outside of the auditory cortex in order to support adaptive computations that underlie perception and behavior^64^. These types of questions could be answered by applying our method to neural responses beyond the auditory cortex, testing longer segment durations, and using multimodal stimuli.

Understanding the mechanisms of multiscale temporal processing is an active area of research, and multiple neural mechanisms and computations are thought to play a role. Recurrent connections between neurons, both within and across regions (e.g., feedback), can induce multi-scale temporal processing. For example, recurrent neural networks trained to recognize speech information learn to diversify their integration windows across units and layers, qualitatively analogous to what we observe in the auditory cortex^7^. Multiscale temporal processing can also be induced by synaptic dynamics at the level of individual neurons^65^. Testing the underlying neurobiological mechanisms of multiscale integration is an important topic for future research that will likely involve more detailed neuroanatomical studies combined with computational modeling^65^. This research will benefit from the fact that our TCI method can be applied to any neural or model response, and thus can be used to evaluate whether a given computational model or circuit mechanism can predict the empirically measured integration window.

## Methods

The data from the paper come from three experiments. Experiment I compared integration windows between different regions using natural speech, music, and ferret vocalizations. Experiment II examined layer effects. Experiment III examined the importance of the information rate by comparing original, stretched, and compressed sound stimuli. Experiments I and III were conducted at the École Normale Supérieure (ENS) using chronic multi-electrode arrays recording multi-units, and Experiment II was conducted at Oregon Health & Sciences University (OHSU) using laminar probes recording single units. The neurophysiological recording methods for chronic arrays and laminar probes are described separately.

### Neurophysiological recordings with chronic arrays at ENS

#### Animal preparation

Experiments were performed in adult female ferrets (Mustela putorius furo, ferret A, B, and C) across one or both hemispheres (ferret A: left and right, ferret B: right, and ferret C: left). The animals were 1–3 years of age, weighing 500–1000 g, and were housed in pairs or trios with a normal day-night light cycle and unrestricted access to food and water. All ferrets were virgins. Experiments were approved by the French Ministry of Agriculture (protocol authorization: 21022) and all manipulations strictly comply with the European directives on the protection of animals used for scientific purposes (2010/63/EU).

#### Surgical procedure

Chronic neurophysiological recordings were performed head-fixed after implantation of a metal headpost attached to the skull (detailed procedure in Chillale et al., (2023)^66^). After recovery from the initial headpost surgery, chronic floating multielectrode arrays were implanted in a subsequent surgery. A 10 mm x 10 mm craniotomy was performed under anesthesia (1% isoflurane) over the auditory cortex. The large craniotomy allowed us to identify the different regions of the auditory cortex (anterior/middle/posterior ectosylvian gyri) by visual inspection of the pseudosylvian and suprasylvian sulci and additional landmarks. The animals were then implanted with floating multielectrode arrays (Platinum/lridium, MicroProbes, electrodes of impedance of 500 or 750 kΩ, 4 rows of 8 electrodes with 0.4 μm distance between the electrodes) in primary or non-primary regions using a micromanipulator and sealed with Vetbond (3M). When possible, the dura was pulled over the implanted array. Finally, the removed skull flap was put back and sealed with Kwik-Sil (WPI) and bone cement (Palacos, Heraeus). Animals could then recover for 1 week, with unrestricted access to food, water, and environmental enrichment.

#### Recording procedure

Recordings were performed head-fixed in a soundproof chamber. Continuous electrophysiological recordings were digitized, amplified, and recorded at 30 kHz using a digital acquisition system (OpenEphys). We collected 19 recording sessions in Ferret A, 45 in Ferret B, and 40 in Ferret C.

#### Functional localisation of the multielectrode arrays

Floating microelectrode arrays (FMAs) spanned a 2×4 mm cortical area and were implanted following large craniotomies (>1 cm²). Implantation was guided by clearly visible anatomical landmarks on the cortical surface, specifically the suprasylvian (sss) and pseudosylvian (pss) sulci. Arrays placed ventral to the pss targeted non-primary auditory areas, whereas those positioned medio-posterior to the pss were within primary regions (AEG). The recording locations were further verified physiologically through established dorsal-to-ventral tonotopic gradients, confirmed across animals, and consistent with previously published auditory cortical maps^16,45,67^. Thus, both anatomical landmarks and physiological measures reliably determined cortical recording sites. Functional response properties, particularly frequency tuning gradients, provide an independent confirmation of anatomical localization. The convergence of both anatomical landmarks and physiological measures ensured accurate determination of cortical recording sites. All of our anatomical findings are based on large-scale, region-level differences that are robust to any small localization errors that may exist. To verify robustness, we repeated our key anatomical analyses (e.g., comparing MEG and PEG), excluding cells that fell near the boundary between regions.

#### Preprocessing

Electrode responses were common-averaged to the grand mean across all functional electrodes of each array. We applied a 300-6000 Hz band-pass filter (2^nd^ order Butterworth filter with 3dB cutoff applied forward and backward). We then detected multi-unit activity by thresholding the signal (3 SD) and removing artifacts using PCA-based customized spike sorting routines written in MATLAB^68^.

### Neurophysiological recordings with acute laminar probes at OHSU

#### Animal preparation and surgical procedure

Adult male ferrets (aged 6–9 months) were surgically implanted with a head post to stabilize the head and enable multiple small craniotomies for acute electrophysiological recordings. Anesthesia was induced with ketamine (35 mg/kg) and xylazine (5 mg/kg) and maintained with isoflurane (0.5–2%) during the surgical procedure. The skin and muscle on top of the cranium were removed, and the surface of the skull was cleaned. Ten to twelve small surgical screws were placed on the edge of the exposed skull as anchor points. The surface of the skull was chemically etched (Optibond Universal, Kerr), and a thin layer of UV-cured dental cement (Charisma Classic, Kultzer) was applied over the exposed surface. Two stainless steel head posts were aligned along the midline and embedded with additional cement. Finally, cement was used to build a rim extending out from the edges of the implant. The rim served the dual purpose of holding bandages over the implant margin wounds and creating wells to hold saline over the recording sites. Once the implant was finished, excess skin around it was removed, the wound around the implant was closed with sutures, and the animal was bandaged. Antibiotics and analgesics were administered as part of the post-op recovery.

After at least 2 weeks following surgery, the animals were acclimated to a head-fixed posture during intervals starting at 5 min and increasing by 5–10 min every day. Food and liquid rewards were given during these acclimation sessions to help the animals relax under restraint. Animals were considered ready for recording when they could be restrained for more than 3 h without signs of distress (e.g., the animals being relaxed enough to fall asleep).

#### Functional localisation of the single units

The putative location of A1 and dPEG was determined during the headpost implantation surgery based on external landmarks: the posterior and medial edges of A1 falling, respectively, 13 mm anterior to the occipital crest and 8 mm lateral to the center line, and dPEG immediately antero-lateral to A1^45^. To functionally confirm recording locations, we opened small craniotomies <1 mm diameter and performed preliminary mapping with tungsten electrodes (FH–Co.. Electrodes, AM Systems Amp, MANTA software^69^). We measured the tuning of the recording regions using rapid sequences of 100 ms pure tones and used tonotopy to identify cortical fields. We specifically looked for the frequency tuning inversion: high-low-high moving in an antero-lateral direction, which marks the boundary between primary (A1) and secondary (dPEG) fields.

#### Preprocessing

At tonotopically mapped sites, we performed acute recordings with 64-channel integrated UCLA probes^70^, digital head-stages (RHD 128-Channel, Intan technologies), and OpenEphys data acquisition boxes and software^71^. The probes were inserted approximately normal to the cortical surface, up to a depth of ∼1 mm from the dura surface. Due to spatial constraints of the recording site and apparatus, penetrations deviated from normality up to 20°. The depth of recording sites and their location in superficial areas A1 or PEG were confirmed by current source density analysis. Raw voltage traces were processed with Kilosort 2^72^, clustering and assigning spikes to putative single neurons. The clusters were manually curated with Phy^73^. Units were only kept for analysis if they maintained isolation and a stable firing rate over the course of the experiment. Unit isolation was quantified as the percent overlap of the spike waveform distribution with neighboring units and baseline activity. Isolation >95% was considered a single unit and kept for analysis. We further filtered neurons based on the reliability of their responses, requiring a Pearson’s correlation >0.1 between PSTH responses to the same stimuli (3-10 repetitions, 20 Hz sampling) drawn from random halves of repeated trials.

### Experimental protocols

#### Stimuli

The sound sequences in all experiments were created using segments of varying durations excerpted from natural sounds (examples in **Supplementary** Figure 8). Each segment duration was created by dividing longer segments into as many shorter, consecutive segments as each duration allowed. For example, 250 ms segments were created by subdividing the 500 ms segments into two segments at the midpoint. The same was done to create 125 ms segments from the 250 ms segments. Segments were slightly longer than the target duration (by 15.625 ms) in order to enable cross-fading between the segments. In Experiments I and III, the sounds were sampled at 44.1kHz, amplified (Sennheiser HDVD 800), and presented via custom scripts implemented using PsychToolbox (MATLAB version 3.0.18^74–76^) through a central free-field speaker (Visaton SC 8 N - 8 Ohm 70-20,000Hz). In Experiment II, the digital acoustic signals were transformed to analog (National Instruments), amplified (Crown D-75A), and delivered through free-field speakers (Manger W05, 50–35,000 Hz flat gain). Sounds were presented 30 deg azimuth controlatéral from the recording site (0 ° elevation; 60 dB SPL).

#### Experiment I

The goal of this experiment was to measure integration windows using a diverse set of natural sounds. The stimuli tested included sequences composed of segments of varying durations (ranging from 15.625 to 500 ms). Segments were excerpted from 24 natural sounds (each 500 ms) spanning three different categories (8 sounds/category): speech, ferret vocalizations, and music. Each sound was subdivided to create segments of the desired duration. Each natural sound was RMS-normalized prior to segmentation. We presented the segments in two pseudorandom sequences, with all segment durations and categories interleaved. Sequences varied in duration from 2 to 6 minutes, depending on the session and animal. Segments were concatenated using cross-fading to avoid click artifacts (15.625 ms raised cosine window). Each stimulus was repeated several times to measure a reliable noise ceiling (8 repetitions for most recording sessions).

#### Experiment II

The goal of this experiment was to replicate key findings from Experiment I as well as characterize the organization of integration windows across cortical layers using acute laminar probes. The stimuli tested were similar to those in Experiment I. The segments were excerpted from 20 natural sound recordings that included ferret vocalizations, speech, music, and other animal calls/vocalizations and environmental sounds. The segment durations spanned 16 to 500 milliseconds. The segments for each duration were presented in a separate sequence, using two different random orders.

#### Experiment III

The goal of the experiment was to examine whether integration windows reflected absolute time or the rate at which information varies in sound. Stimuli were similar to those for Experiment I except that we only tested speech and ferret vocalizations (not music), and we altered the information rate of the sounds by stretching and compressing the sounds (without altering the pitch).

Sounds were stretched and compressed by a factor of and prior to subdivision into segments using a phase vocoder with identity phase locking^77,78^. Segments were excerpted from 16 natural sounds (500 ms, half speech, half ferret vocalizations) using the same range of segment durations as Experiment I. We also tested natural rate sounds without any stretching or compression.

### Summary of replications

There are 5 key results from our study: (1) Neurons show a time-limited integration window beyond which stimuli have very little effect on the neural response. (2) Integration windows are diverse, varying by approximately 3 octaves across the population. (3) Integration windows increase substantially between hierarchically organized regions, specifically MEG and PEG. (4) Integration windows do not differ substantially between cortical layers. (5) Integration windows do not vary with the information rate of sound.

Results 1 and 2 are replicated across all three experiments. Result 3 is replicated across experiments 1 and 2, and could not be tested in Experiment III due to insufficient data from the primary auditory cortex. Result 4 was only tested in Experiment II because that was the only experiment with laminar probes. Result 5 was only tested in Experiment III. There were no failed replication attempts.

**Table 1.** Summary of the differences between the two experimental sites. Columns represent two experimental sites (OSHU in the US and ENS in France). The rows represent differentiating variables.

|  | US | France |
| --- | --- | --- |
| Age | 6-9 months | 1-3 years |
| Sex | male | female |
| Recording type | acute laminar probes | chronic multi-electrode arrays |
| Sound system | free field central speaker; 30° azimuth<br>controlatéral from the recording site | free field central speaker |

### Data analysis

#### Selecting units with a reliable response to sound

Our analyses were performed on unit response timecourses computed by counting spikes within 5 ms bins. This bin width is substantially shorter than the integration windows of units in the auditory cortex and thus unlikely to bias the measured integration window. When plotting the cross-context correlation and noise ceiling (e.g., Figure 1C), we used a larger bin size (10 ms) because we found this aided visualization due to more reliable and less variable measures. Results were similar for 5 and 10 ms bin sizes. We selected units (single and multi-units) with a reliable response time course (binned spike counts) to sound. Specifically, we measured the split-half Pearson correlation of the response timecourse of each unit across all stimuli from two non-overlapping sets of stimulus repetitions (odd vs. even). For Experiments I & II, we selected units with a split-half correlation of at least 0.1, where this correlation was highly significant (p< 10^−5^). We used a lower absolute correlation threshold (0.05) in Experiment III because we were focusing on PEG, which has sparser and more variable responses. We identified 693 units across 8 animals and 3 experiments that showed a reliable response to natural sounds based on these criteria (Exp I: 345, Exp II: 231, and Exp III: 117). Significance was measured using a permutation test where we permuted 100 ms segments of the neural timecourse (1000 permutations) and re-measured the correlation coefficient to build up a null distribution. This null distribution was then fitted with a Gaussian, from which we then computed the tail probability of the observed correlation value. We used a Gaussian fit rather than the empirical distribution so that we could measure small p-values that would be impossible to measure via counting.

#### Cross-context correlation

We measured integration windows using an analysis developed in prior work for estimating context invariance. We estimate context invariance using the cross-context correlation (CCC) and then use a computational model to infer a window that best predicts the CCC. The Results section provides a description of the core logic and analysis of the CCC because it is critical to understanding the results. In order to ensure that the Methods are complete, we reiterate some of that description here, but add more detail and corresponding equations. The analysis is done separately for every unit, and we therefore do not include the unit index in our equations to simplify notation.

The first step of our analysis is to reorganize the response time course of each unit as a matrix. A single element of this matrix, 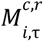, contains the response (binned spike counts) to a segment *i* at time lag τ for context *c* (for a single unit) and repetition *r* (Figure 1B**, Supplementary** Figure 1). Thus, there is a separate matrix for each of the two contexts, and corresponding rows contain response timecourses to corresponding segments. Corresponding columns contain the response to many different segments for a single time lag.

To compute the CCC, we correlate corresponding columns across matrices from different contexts (Figure 1B**, Supplementary** Figure 1A) and average these correlations:

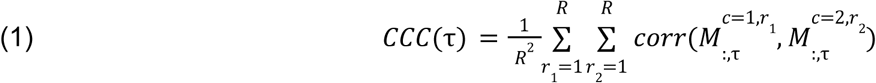

Where 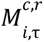 indicates a vector across segments for a single time lag (τ), context (*c*), and repetition (*r*). We use the Pearson correlation. Note that we compute the CCC using all valid pairs of repetitions for different contexts, which is why there is an outer summation across repetitions.

To compute the noise ceiling, we correlate across matrices from different repetitions for the same context and average these correlations across contexts (**Supplementary** Figure 1B):

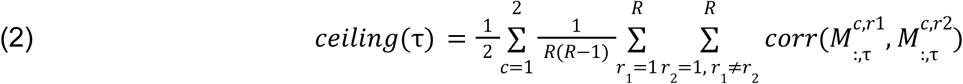

We found that the noise ceiling was, on average, similar across different cortical regions, layers, sound categories, and sound rates (**Supplementary** Figure 9A-D). We found a weak negative correlation between the integration width and the noise ceiling across units (**Supplementary** Figure 9E)(ρ = −0.32, p<0.001), with lonshorter ger integration windows tending to have higher noise ceiling. Importantly, the noise ceiling is computed separately for each unit and stimulus/time-lag, and thus accounts for any variation in the response reliability across units and stimuli. In our prior work, we have shown using simulations that our method can reliably estimate integration windows across many different signal-to-noise ratios^30^.

Shorter segments were embedded within longer ones; we could therefore define two distinct context comparisons: (1) random context, where a segment is surrounded by unrelated segments, and (2) natural context, where the shared segment is a part of a longer one, preserving its original temporal surroundings (visual schematic Figure 5A). To ensure meaningful comparison, the two contexts must differ, requiring at least one to be random. The second context could be either random or natural. In our analyses, comparisons between two random contexts and those between a random and a natural context yielded comparable results (Figure 5B-D). To maximize statistical power, we computed the cross-context correlation by pooling comparisons from both context types.

We note that the cross-context correlation and noise ceiling are computed across segments. Since there are fewer segments for the longer segment durations, our measures will tend to be more variable for longer segments.

#### Model-estimated windows

In our prior work, we developed a computational model that is capable of using the cross-context correlation from multiple segment durations and lags to estimate a single underlying integration window^30^. We have extensively tested and validated the method in this prior work, showing that we can recover accurate integration window estimates from multiple ground truth models using noisy data.

The integration window is parametrized by a Gamma distribution, which is a simple, causal distribution that is commonly used to model temporal windows. We begin by defining a single-parameter Gamma distribution (*g*) that varies in its shape (β) from more Exponential to more Gaussian:

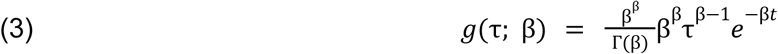

This window(ℎ) is then shifted and scaled to accommodate different widths and centers:

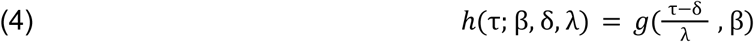

The shift and scale parameters (δ, λ) do not correspond directly to the width (*w*) and center (*c*) because the scale parameter (λ) alters both the width and center. These parameters were instead computed from the width and center as follows:

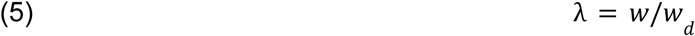

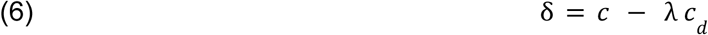

In these equations *w_d_*, and *c_d_* are the width and center for our baseline Gamma distribution without any scaling or shifting (i.e., *g*(τ, β) = ℎ(τ, β, δ = 0, λ = 1). The baseline center (*c_d_*) was calculated using the inverse cumulative distribution function of this baseline distribution. The width (*w_d_*) does not have a simple mathematical form that we are aware of. It was therefore calculated empirically by exhaustively measuring the probability mass of all possible intervals of the baseline distribution (using the cumulative distribution function), and selecting the smallest interval that contains 75% of the mass (with a tolerance of 10^-6^).

The cross-context correlation will depend on the extent to which the integration window overlaps the shared segment vs. the surrounding segments. We therefore start by measuring the overlap of the model window with each segment. The overlap varies with the time-lag between the window and the segment and was computed by convolving the window with a boxcar function (*b*(*t*)) with a 1 for all timepoints where the segment was present:

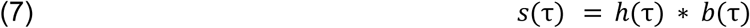

To account for cross-fading, the edges of the boxcar were tapered using the same Hanning window used for cross-fading (≈7.8 ms ramp). We then computed a prediction for the cross-context correlation (*P_CCC_*(τ)) by multiplying the noise ceiling by a ratio that reflects the relative overlap of the shared

(*S_shared_*(τ)) vs. surrounding (*S_n,surround_*(τ)) segments:

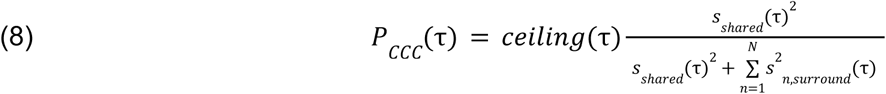

This functional form was derived in our prior paper and has been empirically validated through extensive simulations^30^.

We determine the best fitting window by varying the parameters of the window and selecting the window that yields the best prediction of the cross-context correlation (width spanning 100 logarithmically spaced widths between 1/128 to 0.5 seconds, center spanning 100 logarithmically spaced centers between 0 to 0.25 seconds, shape=[1,2,3,4,5]; parameter combinations that led to an acausal window were excluded).

#### Relationship to standard tuning parameters

We investigated the relationship between integration width and three standard tuning parameters: best frequency, bandwidth, and evoked spike rate (**Supplementary** Figure 10). We used a hierarchical Linear Mixed Effects model to examine the effect of these parameters, including distance to primary auditory cortex as a covariate to account for effects of the anatomical hierarchy. In practice, we found that bandwidth was highly correlated with best frequency (*r* = 0.9, *p* < 0.001), and we only included best frequency in the model. We observed significant effects of best frequency (F_1,340_ = 4.01, *p* = 0.045, β_best frequency_ = −0.103 octaves/octave, CI = [−0.204, −0.002]) and evoked spike rate (F_1,342_ = 16.08, *p* < 0.001, β_evoked spike rate_ = 1.7 octave per spike/sec, CI = [0.866, 2.533]; N = 345). However, the effect sizes for these variables (beta weights for standardized predictions (β_best frequency_ = −0.059, β_evoked spike rate_ = 0.106) were much smaller (∼3-6x smaller) than that for distance to primary auditory cortex (β_distance to PAC_ = 0.327) (best frequency: F_1,334.86_ = 146.91, *p* < 0.001; spike rate: F_1,337.45_ = 28.96, *p* < 0.001).

#### Anatomical analysis - cortical surface maps

There is variability in the functional location of cortical fields across animals. To create a cortical surface representation of all recorded units and to account for functional variability, we localized each electrode array on a standardized template tonotopic map, computed across many ferrets^79,80^. The location of each array was selected based on the frequency selectivity of the electrode responses within that array, the tonotopic gradient found across the array, and its position relative to anatomical landmarks. Electrode recordings from different sessions were assumed to sample different cells, and each recording was therefore plotted at the same location with a small amount of jitter so they could be distinguished. The jitter was sampled using polar coordinates (radius was sampled from a uniform distribution from 0 to 360°, and distance was sampled from a uniform distribution from 20 to 240 micrometers). Recordings from the left hemisphere were mirrored onto the right hemisphere for visualization purposes. We measured the distance on the cortical surface of each unit to a reference point in the middle of the primary auditory cortex.

#### Anatomical analysis - laminar assignment

Laminar analysis was conducted following a method presented in our previous paper^81^. In summary, cortical layers were categorized into three groups (layers 1–3, layer 4, and layers 5–6) and assigned to each electrode of the silicon probes using a custom graphical user interface. Boundaries between these groups were determined based on characteristic features of the local field potential (LFP) signal. The LFP signal was extracted by applying a zero-phase shift, 4th-order Butterworth low-pass filter (at 250 Hz) to the raw electrophysiological data, followed by downsampling to 500 Hz. Key features used for layer identification included the current source density (CSD) sink and source patterns elicited by broadband noise bursts centered on each site’s best frequency (BF), which align with established auditory-evoked CSD patterns in the auditory cortex of other species^82^. Additionally, the relative power of high-frequency (>40 Hz) versus low-frequency (<30 Hz) components of the LFP was analyzed to delineate layers 1–3, leveraging prior findings of increased high-frequency power in superficial cortical layers^83^. Finally, layer 4 was identified by detecting a drop in LFP coherence between adjacent channels at its upper boundary, consistent with earlier reports^84,85^. Each unit was assigned to a cortical layer based on the electrode exhibiting the highest spike amplitude for that unit.

#### Rate-yoking index

Compressing and stretching change the rate at which information varies by a known magnitude. We used a stretching and compression factor of such that there is an octave difference in the rate at which information varies between stretched and compressed sounds. Thus, if the integration window reflects the information rate, we should observe a one-octave change in the integration window. We therefore computed a rate-yoking index by computing the difference in integration windows on an octave scale between stretched and compressed speech and dividing this difference by that which would be expected for a rate-yoked window (i.e., a one-octave change):

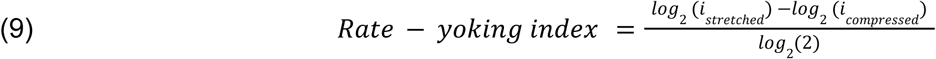

Where *i* is the measured integration window for stretched/compressed sounds. An index of 0 indicates a fixed, time-yoked window, while a value of 1 indicates a fully rate-yoked window. Note that the denominator in the above equation happens to be 1, and thus has no effect on the result, since there was an octave difference in the information rate for stretched and compressed sounds. If we had used a different stretching and compression factor, then it would be different and would need to be included. The rate-yoking index is not strictly bound between 0 and 1, and noise will tend to cause the estimate to exceed these bounds. A value less than 0 indicates that the integration window was longer for compressed sounds, and a value greater than 1 indicates that the difference between stretched and compressed integration windows exceeded 1 octave.

#### Statistics

Unless otherwise noted, we used a linear mixed effects (LME) model to evaluate the significance of our effects (fitlme.m and coefTest.m in MATLAB; Satterthwaite approximation was used to estimate degrees of freedom when measuring significance). Whenever possible, we included random effects to account for variation across sessions from day to day, using diagonal covariance matrices to prevent overfitting/singular results. As is typical, we do not include random effects for animals because of the small number of animals in our experiment. For categorical variables, the category with the most samples was always used as the baseline and not modeled explicitly in the linear model. Integration window estimates were always logarithmically transformed before being fit by the LME model.

For Experiment I, to test whether integration windows increased across the cortical hierarchy, we modeled the integration window as a linear function of distance to primary auditory cortex, including random intercepts and slopes for different recording sessions:

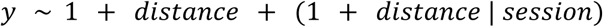

Where *y* reflects either the integration width or center (logarithmically transformed). The sample dimension was unit (single or multi-unit).

To test whether integration windows varied across regions, we used the following model:

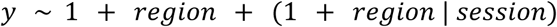

Region was a binary categorical variable (primary vs. non-primary auditory cortex).

For Experiment II, which used laminar probes, we investigated the effects of layer and region using the following model:

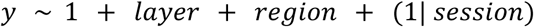

Layer was a categorical variable with three levels (granular, infragranular, supragranular). Region was again a binary variable (primary and non-primary auditory cortex). A different insertion was used for each recording session, and as a consequence, the session was partially confounded with region and layer, and we found that a model with random slopes did not converge (and these terms were thus removed). We tested whether the effect of region was significantly different (larger) than the effect of layer by directly comparing their coefficients against a null model that assumed no difference. To make this a fair comparison, we collapsed our layers from three levels to two levels to make them comparable to those for each region. To be conservative, we chose the two levels so that the effect of layers would be maximal, thus minimizing the likelihood that the effect of the region would be even larger than the effect of the layer. To this end, we separated out the infragranular layers from the granular and supragranular layers (infragranular vs. granular/supragranular), because the infragranular layers showed the most different integration windows (Figure 3).

For Experiment III, we used the following model to investigate the effect of rate, controlling for any possible difference due to sound category (ferrets vs. speech):

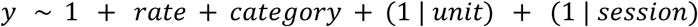

Rate and category were categorical variables with three (original, compressed, speech) and two (speech, ferrets) levels, respectively. We did not include random slopes for sessions in this model because the model was not able to estimate them (variance terms for these random effects were either undefined or close to 0).

To evaluate the effects of tuning parameters on the integration window, we used the following model:

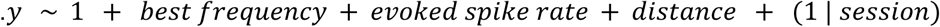

Predictor variables (best frequency, evoked spike rate, and distance to PAC) were standardized by dividing by their standard deviation, such that fixed-effect beta coefficients represent effects in normalized units. For interpretability, slopes and confidence intervals were subsequently back-transformed into the original measurement scales. As in previous analyses, we included random intercepts for recording sessions to account for across-session variability. As for the effect of layer and region, we tested for a significant difference between fixed effects using a Wald F-test (using coefTest.m) with the Satterthwaite approximation for the degrees of freedom. In particular, we tested whether the slopes for best frequency and spike rate differed significantly from the slope for distance. These comparisons were implemented by evaluating the significance of a direct contrast between the variables of interest (e.g., β − β = 0).

### Resource Availability

#### Lead Contact

Requests for further information and resources should be directed to and will be fulfilled by the following contact persons: Magdalena Sabat, Sam Norman-Haignere, and Yves Boubenec.

#### Materials Availability

The analysis code and sample sounds used in the experiments in the paper will be made publicly available upon publication in a peer-reviewed journal.

#### Data Availability

Data necessary for replication of the results in this article will be made publicly available upon publication in a peer-reviewed journal.

## Acknowledgments

We would like to thank Balkis Cadi and Lynda Bourguignon for their technical and administrative support throughout the project. We thank Flavien Feral for his technical support with the surgeries at ENS. This work was supported by the Institut Universitaire de France, ANR-17-EURE-0017, ANR-10-IDEX-0001-02, and ANR-23-NEUC-0001.

## Author contributions

Conceptualization M.S., S.N-H, and Y.B.; Data collection M.S, H.G, Q.G., and M.L.E; Formal Analysis, data curation, and visualization M.S, S.N-H, and Y.B; Software M.S, Q.G., S.N-H, and Y.B; Writing, first draft M.S, S.N-H and Y.B; all authors reviewed the article before submission; Supervision and funding S.N-H, S.V.D, and Y.B.

## Supplementary Figures

**Supplementary Figure 1.**
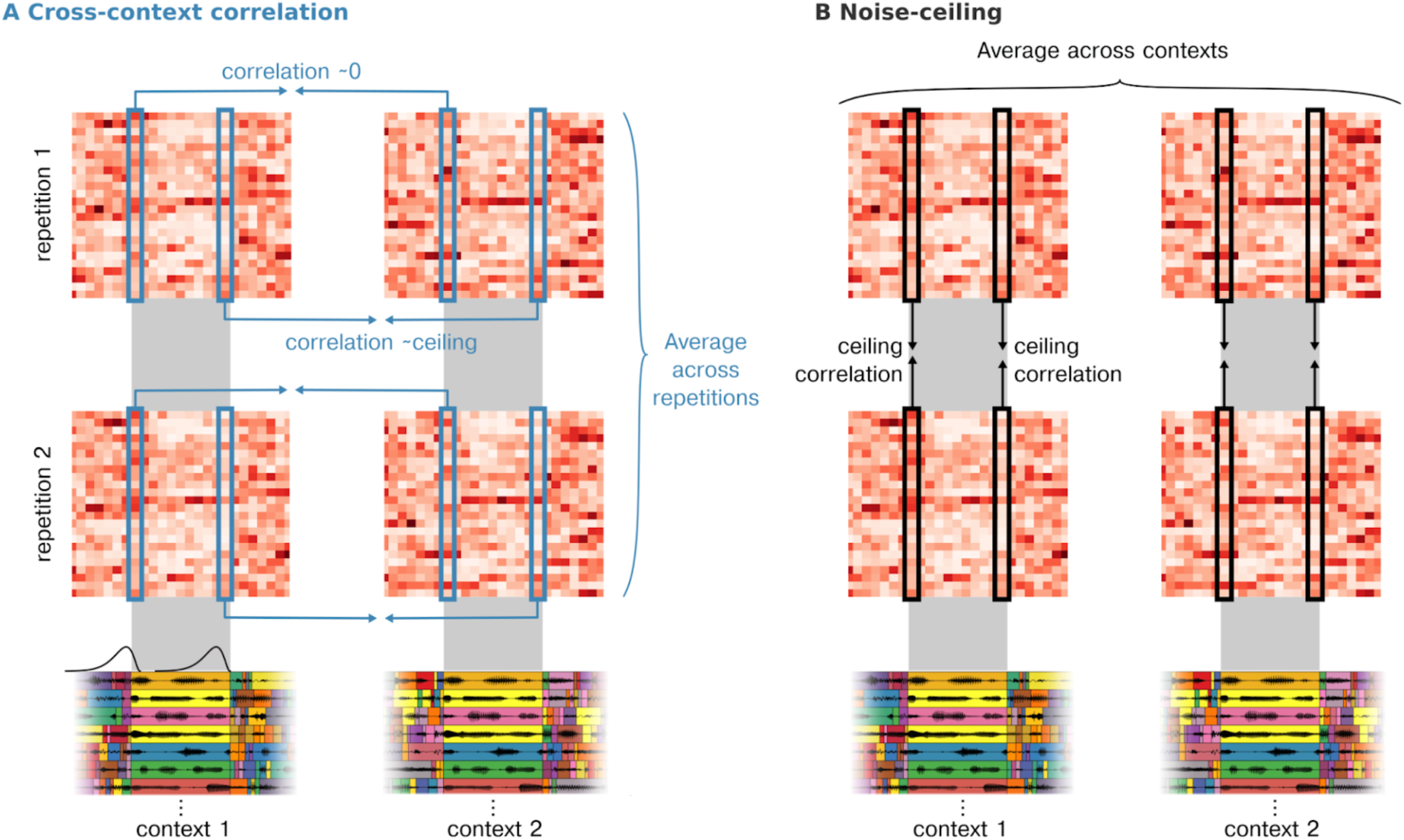
More detailed schematic of the cross-context correlation. (**A**) and noise-ceiling (**B**) as applied to an example unit. This legend gives a brief description of the core ideas as they are linked to this figure. See the text for a more in-depth description of the analyses. **A**, The neural response timecourse of the unit is reorganised as a matrix. Specifically, the response timecourse surrounding all of the segments of a given duration (500 ms in this schematic) was compiled into a segment-by-time matrix, aligned to segment onset (as in **Figure 1B**). The gray shaded area shows the time period when the shared segment was present. Each row contains the response timecourse to a single segment, and each column contains the response to many segments for a single time lag relative to segment onset. Separate matrices were computed for two different contexts (context 1: left, context 2: right) and two different repetitions of the same context (rep 1: top, rep 2: bottom). Below the matrices, we plot waveforms of corresponding segments: the top waveform corresponds to the segment for the first row of the matrix, the second waveform from the top corresponds to the segment of the second row, and so on. The cross-context correlation is computed by correlating corresponding columns of matrices from different contexts (blue columnar boxes), separately for each repetition, and then averaging the correlation coefficients across repetitions. The correlation is computed separately for each time lag relative to segment onset, and a schematic of the hypothesized integration window at each time lag is shown below, overlaid on the stimulus waveforms. At segment onset, the integration window will fall on the preceding context segments, which are independent across contexts, and the correlation should thus be approximately 0. If the integration window is less than the segment duration, then there will be a moment when the window is fully contained within the shared segment, and at this moment, the cross-context correlation should equal the maximum possible value given by the noise ceiling. One can visually observe that the responses from this unit become more similar as time progresses within the segment. To make it possible to visually observe the key trends in this figure, we used a 50 ms bin. Quantitative analyses were performed using a much smaller bin (5 ms) to ensure that the bin size did not upward-bias the measured integration window. **B**, To compute the noise ceiling, we correlated columns across repetitions, separately for each context, and averaged the correlation coefficient across contexts. Because the context is identical, the noise ceiling provides an upper bound of the maximum possible correlation that could be observed when comparing responses across different contexts.

**Supplementary Figure 2.**
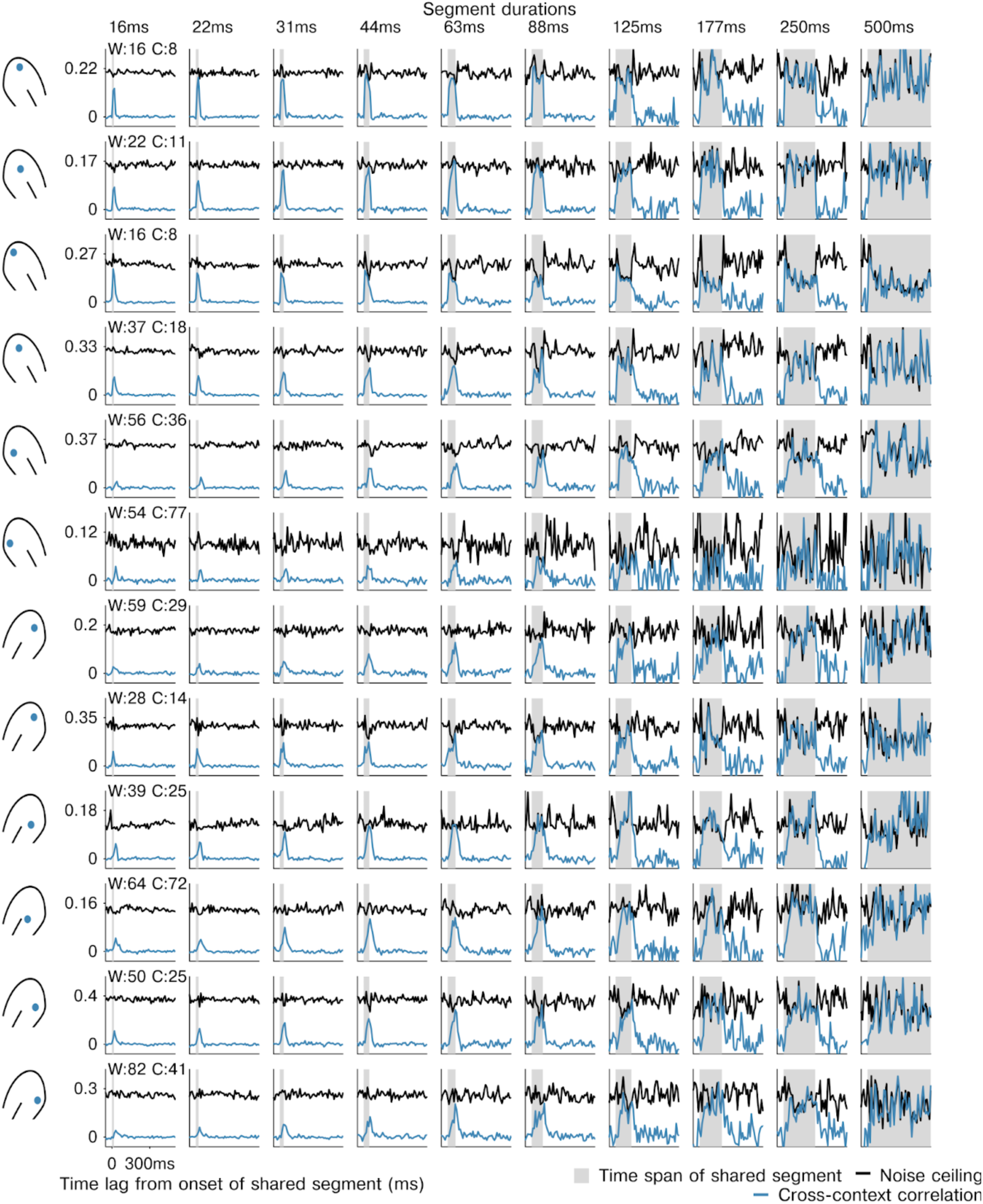
**Examples of cross-context correlation for multi-units in different regions of the auditory cortex**. Examples of the CCC (blue) and noise ceiling (black) from example multi-units from primary and non-primary ferret auditory cortex. For all units, there is a lag and segment duration for which CCC equals the noise ceiling, indicating a context-invariant response. The segment duration needed to achieve a context-invariant response varies substantially across units. The location of each recorded multi-unit is indicated to the left of each row.

**Supplementary Figure 3.**
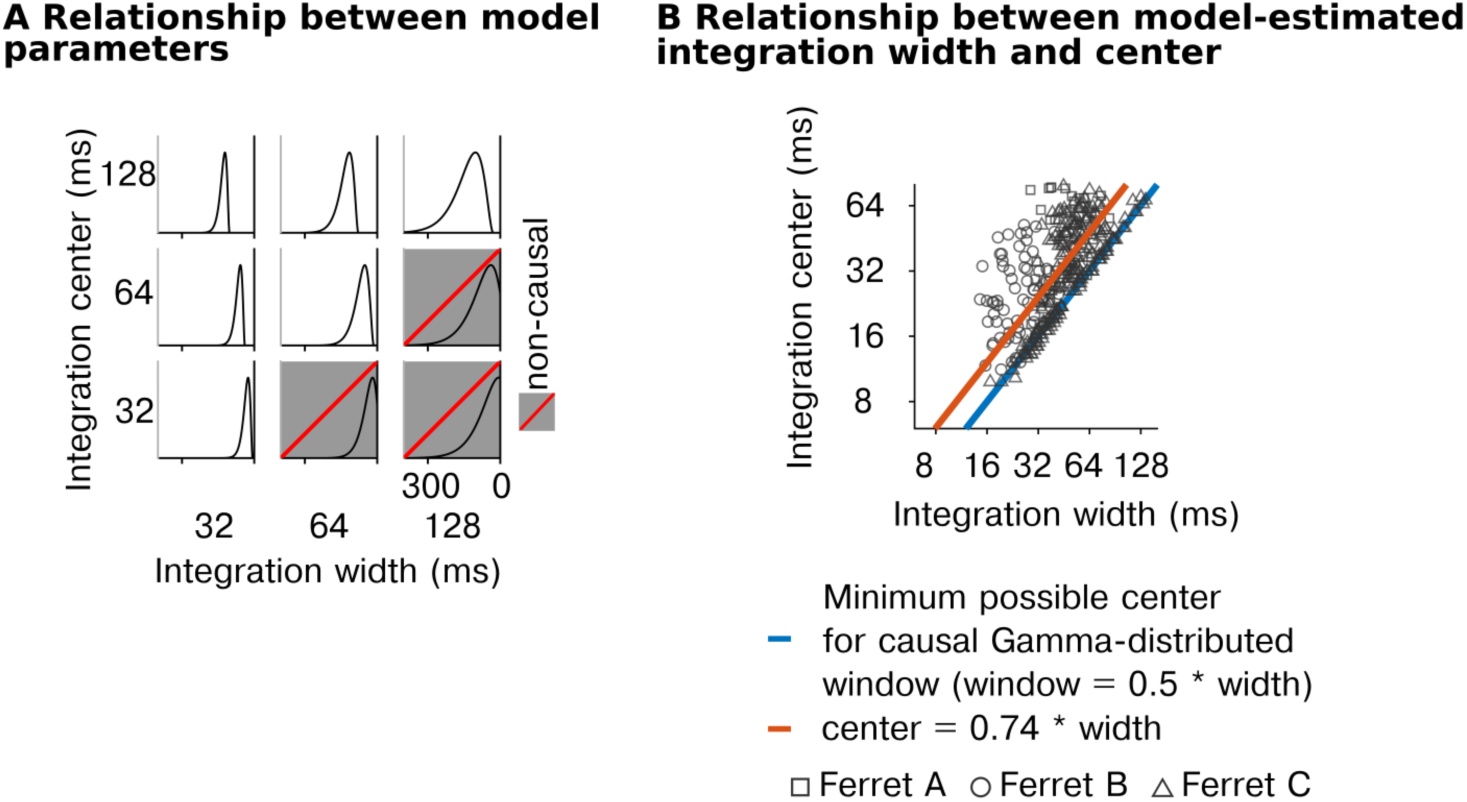
Relationship between the center and width of the integration window. **A**, Integration windows were estimated using a parametric window (Gamma distribution) with a varying width and center. We estimated the width and center that best predicted the cross-context correlation, excluding parameters that yielded a non-causal window. This panel plots examples of the parameter window as a function of the window’s width (x-axis) and center (y-axis). Combinations of parameters that led to acausal windows were excluded (gray box with red dashed line) because they are not biologically possible. **B**, Scatter plot of estimated integration centers vs widths for all units with a best-fit linear function overlaid (orange line). The blue line shows the minimum possible center for a causal, Gamma-distributed window.

**Supplementary Figure 4.**
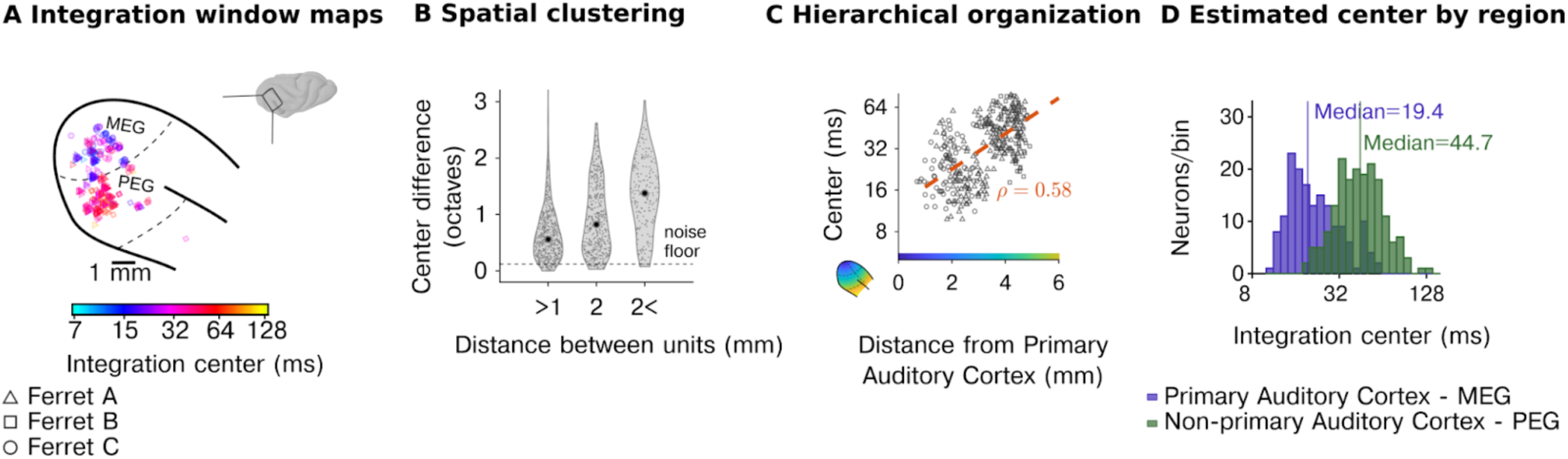
Organisation of integration centers across neural populations in the auditory cortex. **A**, Anatomical map of model-estimated integration centers in three animals (window center: median of the interval containing 75% of the window’s mass). **B**, The difference between the centers of the integration windows between pairs of units as a function of their spatial distance, demonstrating that nearby units have more similar centers. **C**, Centers of the integration windows as a function of distance to the primary auditory cortex (see color map in inset). **D**, Histograms of the centers of the integration windows for primary and non-primary auditory cortex showing substantial diversity across units and hierarchical organization.

**Supplementary Figure 5.**
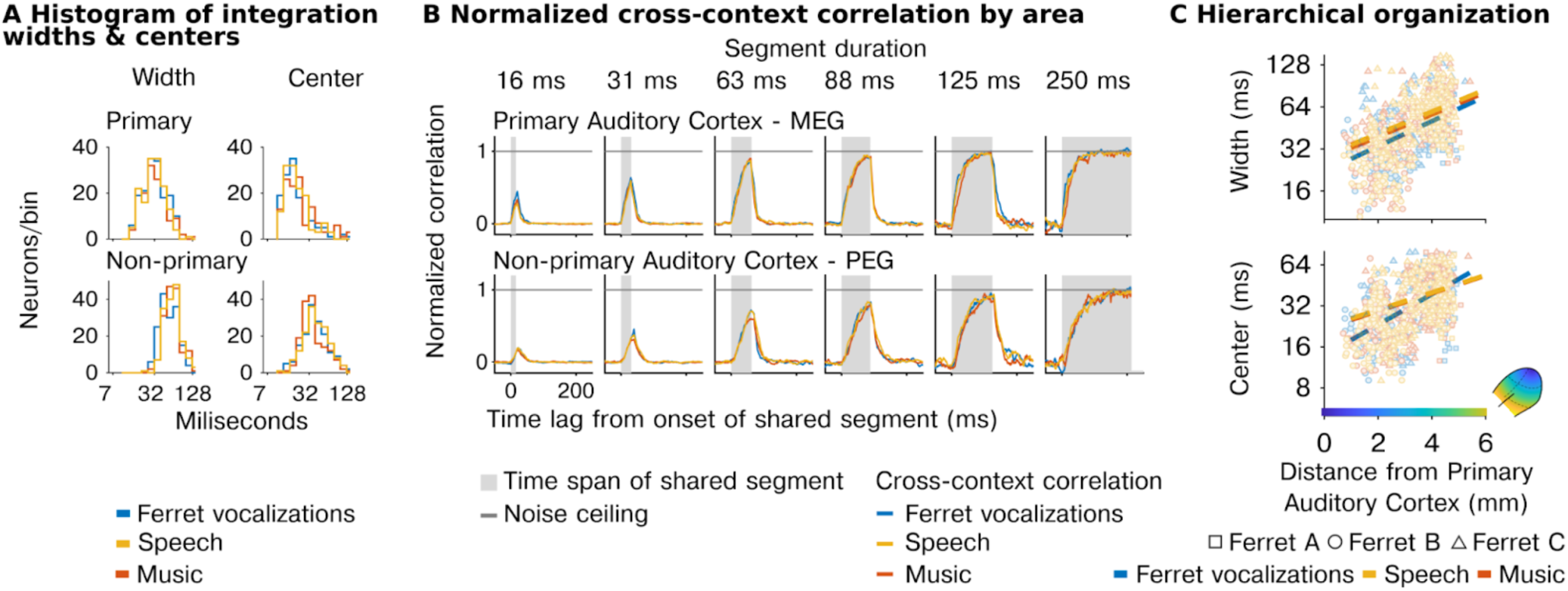
Temporal integration is similar for different stimulus categories. **A**, Histograms of model-fitted integration window parameters (widths and centers) for units in primary and non-primary auditory cortex across the three sound categories tested (ferret vocalizations, speech, music). **B**, Normalized median cross-context correlation for primary and non-primary cortex, plotted separately for each sound category. **C**, Scatter plots of the model-fitted integration centers and widths as a function of distance to primary auditory cortex computed separately for each category with best-fit lines.

**Supplementary Figure 6.**
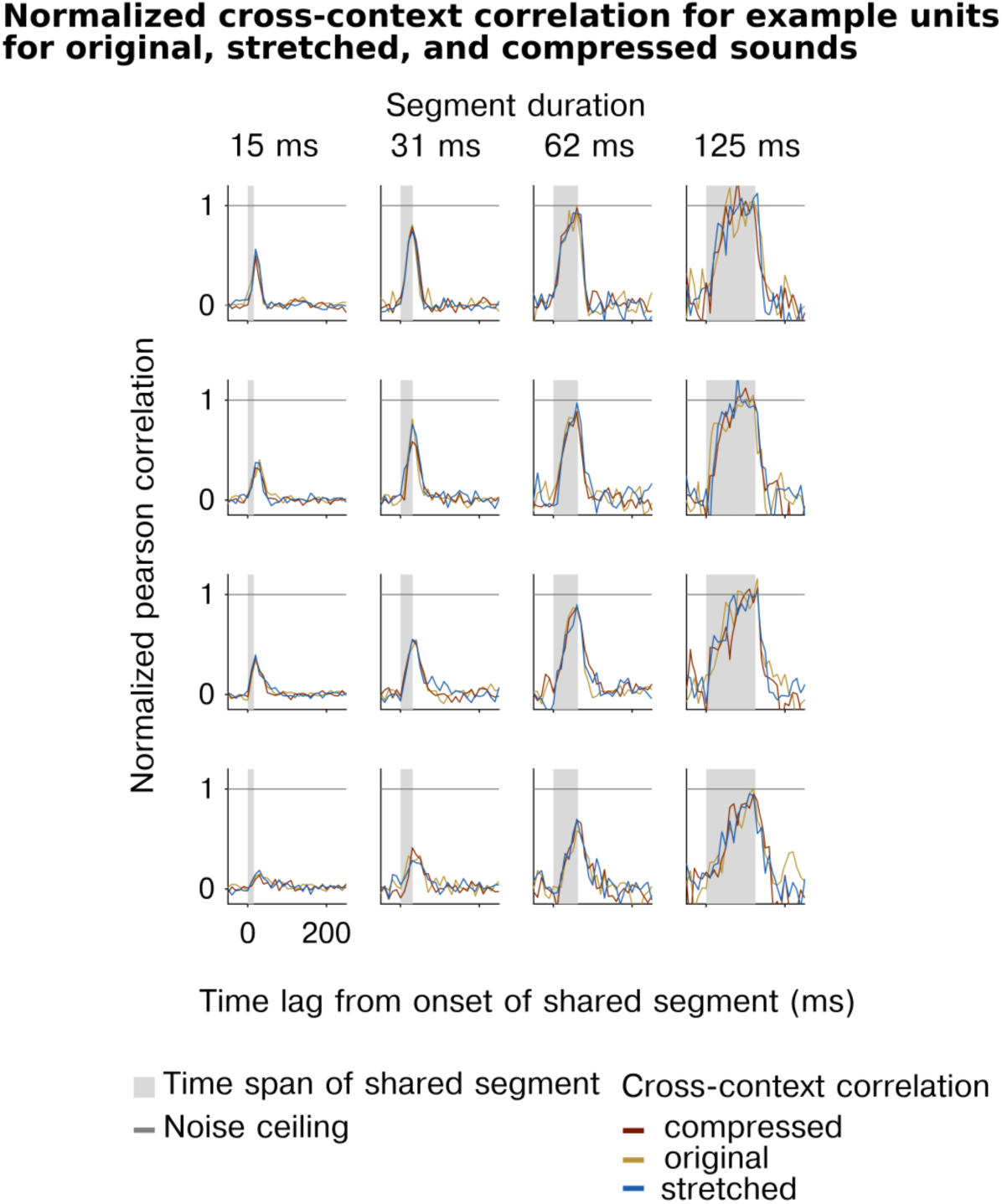
**Normalized cross-context correlation for example multi-units for different sound rates**. Normalized cross-context correlation for five individual units (rows) plotted separately for each of the three different sound rates (compressed, original, stretched). The normalized CCC was computed by dividing the CCC by the noise ceiling separately for every segment duration and time point.

**Supplementary Figure 7.**
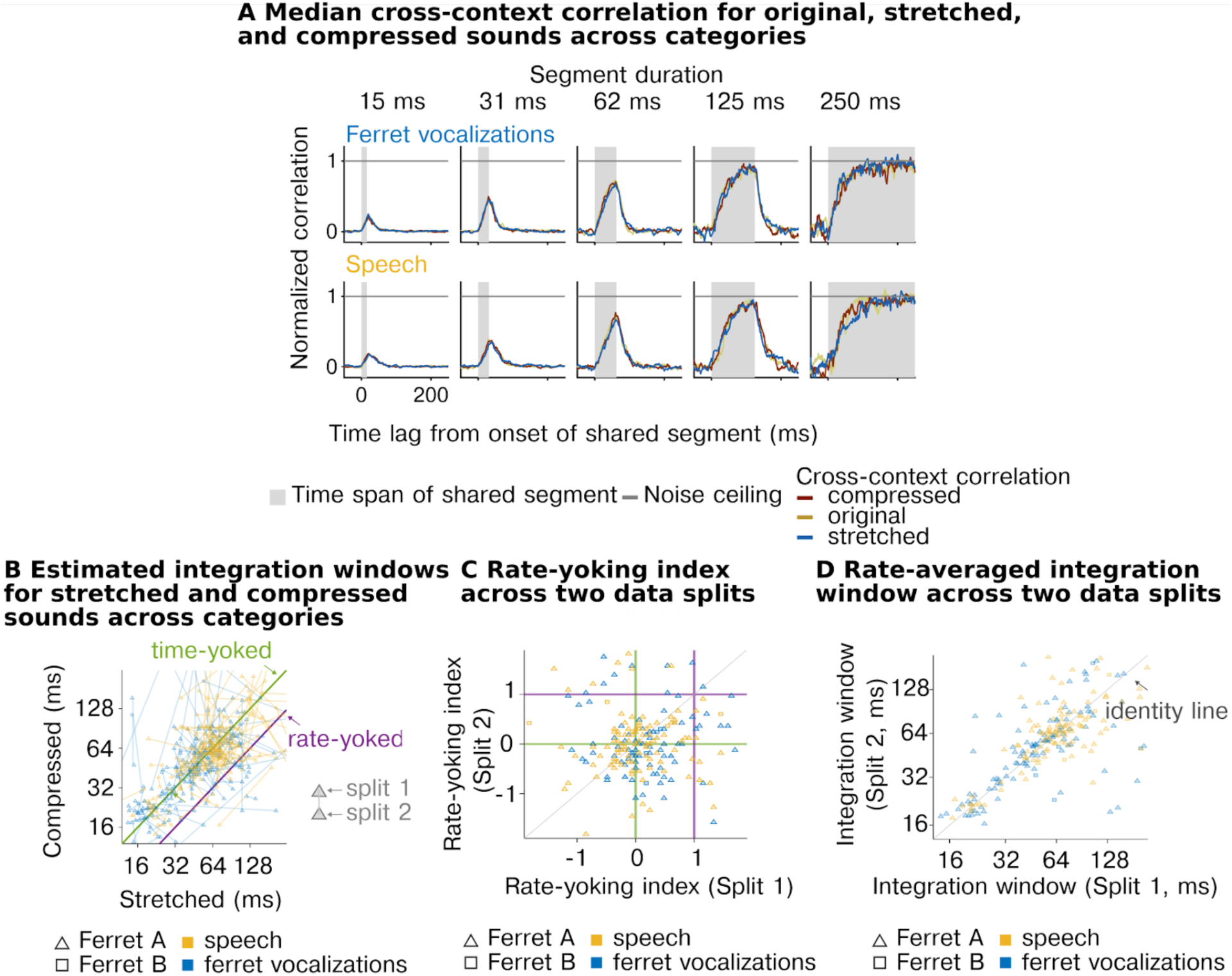
Time-yoked integration is consistent across sound categories. Format is the same as Figure 4, but plotting results separately for speech and ferret vocalisations. **A**, Median normalized CCC across all recorded units for stretched, original, and compressed stimuli computed separately for both sound categories tested (ferret vocalizations and speech). **B**, Integration windows (widths) of all units for compressed (x-axis) and stretched (y-axis) stimuli plotted separately for each sound category (speech in yellow and ferret vocalizations in blue). Green and purple lines show the prediction from a time-yoked vs. rate-yoked response. **C**, Reliability of rate-yoking index across two independent data splits, separately for each category. **D**, Reliability of integration windows averaged across stimuli rates for comparison, again separately for each category.

**Supplementary Figure 8.**
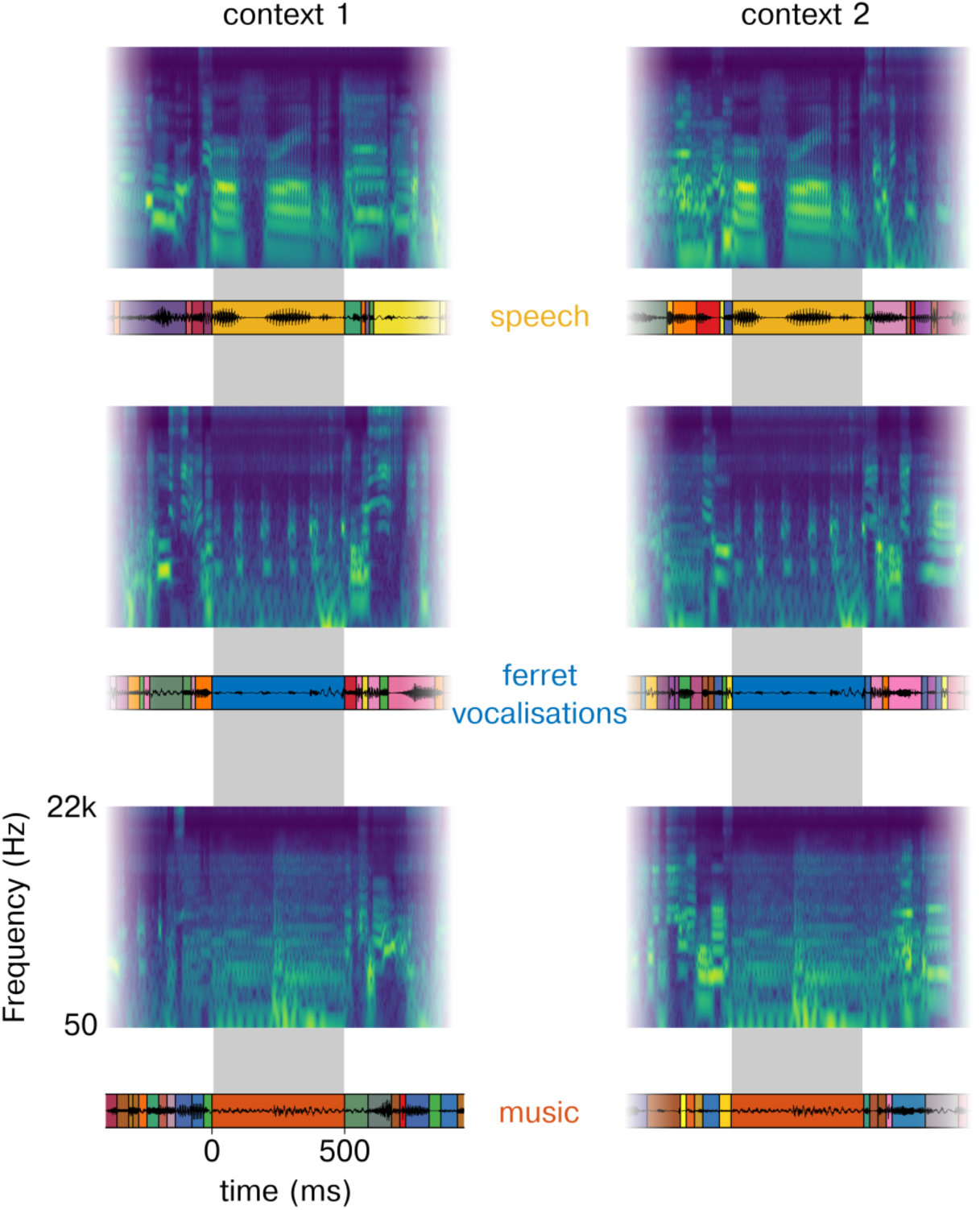
Example spectrograms of the segment sequences used in Experiment I. Each row shows a different example segment (top: speech segment, middle: ferret vocalisation segment: bottom: music segment). The left and right panels show spectrograms of the same segment in two different contexts.

**Supplementary Figure 9.**
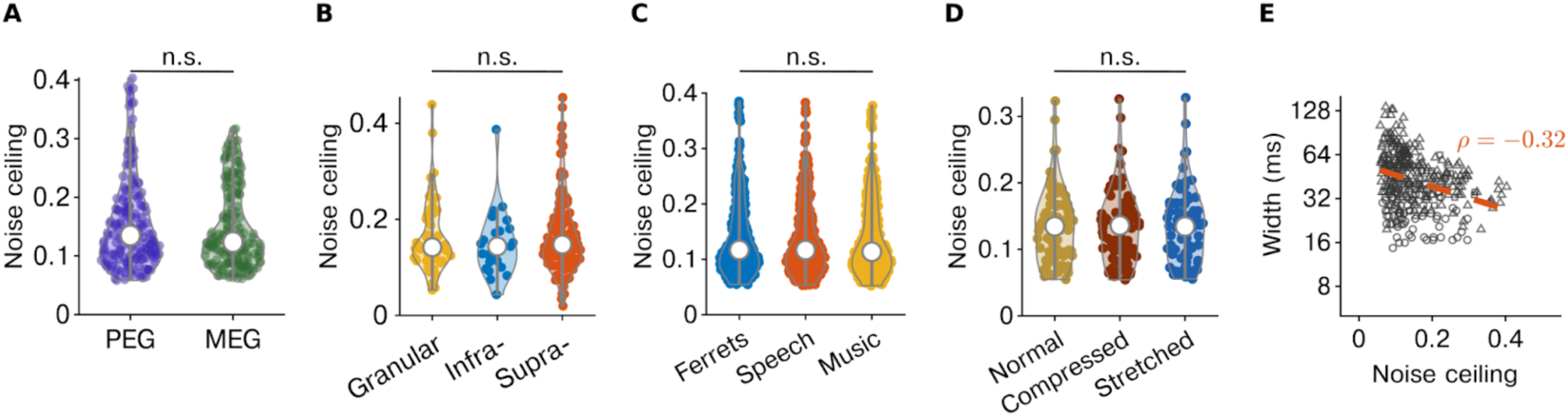
Noise ceiling across conditions and regions. A, Histograms of the distribution of the noise-ceiling in the Primary (PEG) and Non-Primary (MEG) auditory cortex. **B**, Violin plot of the average noise-ceiling for all units recorded in Experiment I across the sound categories. **C**, Violin plot of the average noise-ceiling for all units recorded in Experiment II across the layers. **D**, Violin plot of the average noise-ceiling for all units recorded in Experiment III for the 3 experimental conditions. **E,** Scatter plot representing the relationship between the noise ceiling and integration width (left) and center (right). Orange lines show affine fits, with Spearman correlation coefficients indicated.

**Supplementary Figure 10.**
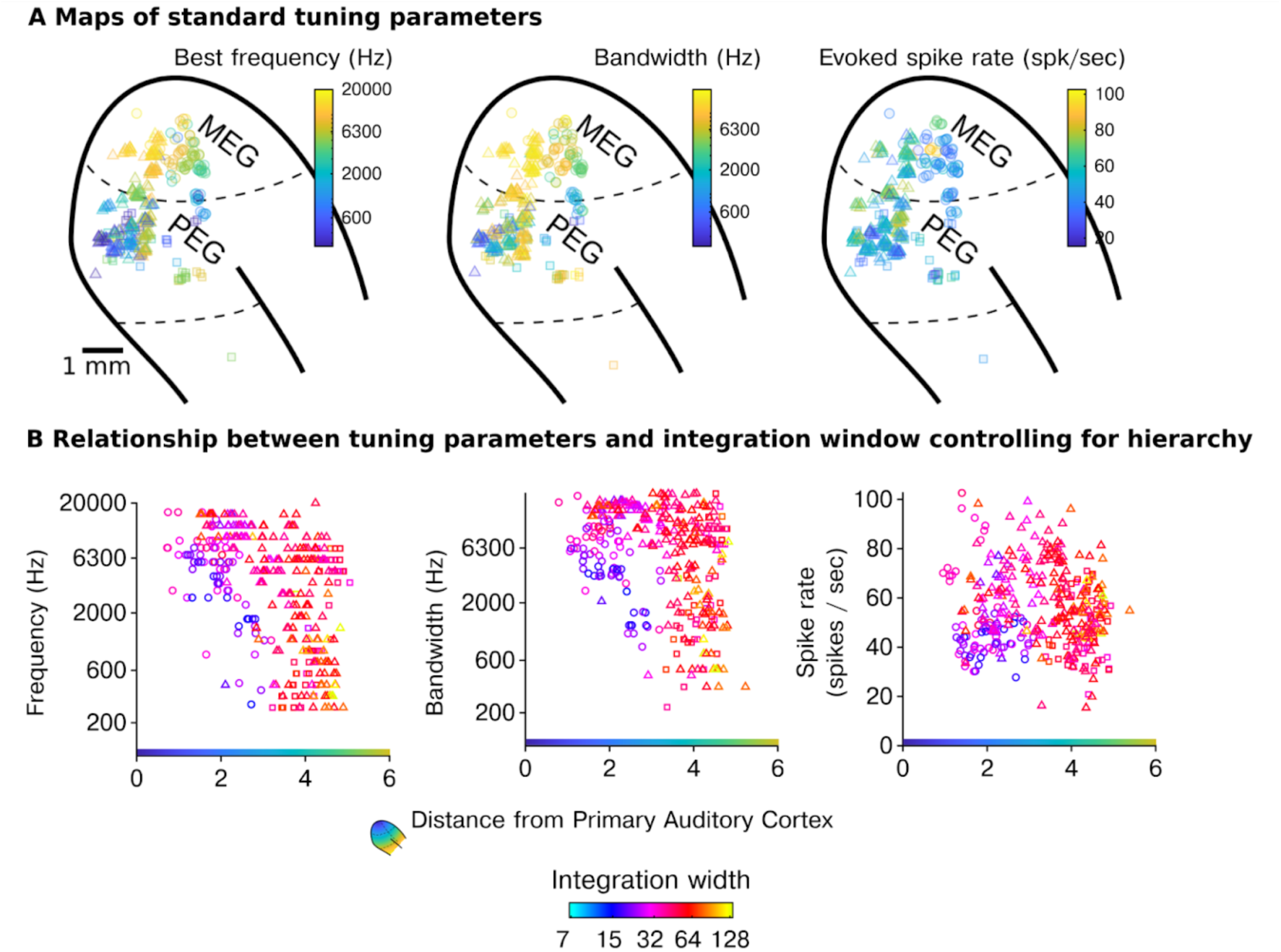
Relationship between standard tuning parameters and neural integration window. A, Maps of standard tuning parameters. **B,** Scatter plots show the tuning parameters for each cell plotted against a measure of anatomical hierarchy (distance to primary auditory cortex) with the integration window indicated by color.

